# White matter microstructure across the adult lifespan: A mixed longitudinal and cross-sectional study using advanced diffusion models and brain-age prediction

**DOI:** 10.1101/2020.04.21.053850

**Authors:** Dani Beck, Ann-Marie de Lange, Ivan I. Maximov, Geneviève Richard, Ole A. Andreassen, Jan E. Nordvik, Lars T. Westlye

**Affiliations:** Department of Psychology, University of Oslo, Oslo, Norway; NORMENT, Division of Mental Health and Addiction, Oslo University Hospital & Institute of Clinical Medicine, University of Oslo, Oslo, Norway; Sunnaas Rehabilitation Hospital HT, Nesodden, Oslo, Norway; Department of Psychiatry, University of Oxford, Warneford Hospital, Oxford, UK; KG Jebsen Centre for Neurodevelopmental Disorders, University of Oslo, Oslo, Norway; CatoSenteret Rehabilitation Center, Son, Norway

**Keywords:** ageing, white matter, multi-shell, longitudinal, diffusion, brain age

## Abstract

The macro- and microstructural architecture of human brain white matter undergoes substantial alterations throughout development and ageing. Most of our understanding of the spatial and temporal characteristics of these lifespan adaptations come from magnetic resonance imaging (MRI), including diffusion MRI (dMRI), which enables visualisation and quantification of brain white matter with unprecedented sensitivity and detail. However, with some notable exceptions, previous studies have relied on cross-sectional designs, limited age ranges, and diffusion tensor imaging (DTI) based on conventional single-shell dMRI. In this mixed cross-sectional and longitudinal study (mean interval: 15.2 months) including 702 multi-shell dMRI datasets, we combined complementary dMRI models to investigate age trajectories in healthy individuals aged 18 to 94 years (57.12% women). Using linear mixed effect models and machine learning based brain age prediction, we assessed the age-dependence of diffusion metrics, and compared the age prediction accuracy of six different diffusion models, including diffusion tensor (DTI) and kurtosis imaging (DKI), neurite orientation dispersion and density imaging (NODDI), restriction spectrum imaging (RSI), spherical mean technique multi-compartment (SMT-mc), and white matter tract integrity (WMTI). The results showed that the age slopes for conventional DTI metrics (fractional anisotropy [FA], mean diffusivity [MD], axial diffusivity [AD], radial diffusivity [RD]) were largely consistent with previous research, and that the highest performing advanced dMRI models showed comparable age prediction accuracy to conventional DTI. Linear mixed effects models and Wilk’s theorem analysis showed that the ‘FA fine’ metric of the RSI model and ‘orientation dispersion’ (OD) metric of the NODDI model showed the highest sensitivity to age. The results indicate that advanced diffusion models (DKI, NODDI, RSI, SMT mc, WMTI) provide sensitive measures of age-related microstructural changes of white matter in the brain that complement and extend the contribution of conventional DTI.

## 1. Introduction

The architecture of human brain white matter undergoes constant remodelling throughout life. Age-related trajectories of white matter macro- and microstructure typically reflect increases in anisotropy and decreases in diffusivity during childhood, adolescence and early adulthood (Krogsrud et al., 2016; Tamnes et al., 2018; Westlye et al., 2010), and subsequent anisotropy decreases and diffusivity increase in adulthood and senescence (Cox et al., 2016; Davis et al., 2009). While the field has primarily been dominated by cross-sectional studies, which by design lack information on individual trajectories (Schaie, 2005), longitudinal studies in the last decade have shown corresponding white matter changes in both development and ageing (Barrick et al., 2010; Bender et al., 2016; Bender & Raz, 2015; de Groot et al., 2016; Likitjaroen et al., 2012; Racine et al., 2019; Sexton et al., 2014; Storsve et al., 2016; Teipel et al., 2010). However, studies that have evaluated individual differences in change across a wide age range are rare (Bender et al., 2016).

White matter properties have commonly been investigated using traditional diffusion tensor imaging (DTI), and the DTI-based metrics fractional anisotropy (FA) as well as mean (MD), axial (AD), and radial (RD) diffusivity are highly sensitive to age (Cox et al., 2016; Sexton et al., 2014; Westlye et al., 2010; Yap et al., 2013). However, limitations of conventional DTI metrics such as their inability to capture restricted non-Gaussian diffusion and lack of specificity to different diffusion pools (Pines et al., 2020) have motivated continued development of more advanced diffusion MRI (dMRI) models. These models include *diffusion kurtosis imaging* (DKI) (Jensen et al., 2005), which was developed to address the restricted diffusion or non-Gaussianity in the diffusion signal; *neurite orientation dispersion and density imaging* (NODDI) (Zhang et al., 2012), which models three types of microstructural environments: intra-cellular, extra-cellular, and an isotropic water pool responsible for the space occupied by cerebrospinal fluid (CSF); *white matter tract integrity* (WMTI) (Chung et al., 2018; Fieremans et al., 2011), which derives microstructural characteristics from intra- and extra-axonal environments; *restriction spectrum imaging* (RSI) (White et al., 2013), which applies linear mixture modelling to resolve a spectrum of length scales while simultaneously acquiring geometric information; and *spherical mean technique multi-compartment* (SMT mc) (Kaden, Kruggel, et al., 2016), a method for microscopic diffusion anisotropy imaging that is unconfounded by effects of fibre crossings and orientation dispersion.

Usually based on multi-shell acquisitions with several diffusion weightings (Andersson & Sotiropoulos, 2015; Jbabdi et al., 2012), these models can be broadly split into “signal” and “tissue” models (D. C. Alexander et al., 2019). Signal representations, such as DTI and DKI, describe the diffusion signal behaviour in a voxel without assumptions about underlying tissue, but as the estimated parameters lack specificity, their characterisation of tissue microstructure remains indirect (Jelescu & Budde, 2017). Tissue models (NODDI, RSI, SMT-mc, and WMTI) involve estimations of the geometry of underlying tissue (Novikov et al., 2019), which may provide higher biological specificity and more precise measures of white matter microstructure and architecture (Jelescu & Budde, 2017; Novikov et al., 2019; Pines et al., 2020). However, despite tissue models being designed to increase specificity, they also require assumptions about the underlying microstructure that may not be fully accurate.

Building on the opportunities from big data in neuroimaging (S. M. Smith & Nichols, 2018), age related brain changes have recently been investigated using machine learning techniques such as brain age prediction; the estimation of the ‘biological’ age of a brain based on neuroimaging data (J. H. Cole et al., 2018; de Lange et al., 2019; Kaufmann et al., 2019; Franke et al., 2010; Richard et al., 2018). Predicting the age of a brain, and subsequently looking at the disparity between predicted and chronological age, can identify important individualised markers of brain integrity that may reveal risk of neurological and/or neuropsychiatric disorders (Kaufmann et al., 2019). While brain age prediction has grown more popular in recent years, most studies have used grey matter features for brain age prediction, while only few have exclusively (Tønnesen et al., 2020), or partly (James H Cole, 2019; Maximov et al., 2020; Richard et al., 2018; S. M. Smith, Elliott, et al., 2019; S. M. Smith, Vidaurre, et al., 2019) utilised dMRI. However, the brain-age prediction accuracy of advanced diffusion models such as RSI and NODDI are not known.

Including cross-sectional and longitudinal data obtained from 573 healthy individuals (with 702 multi-shell dMRI datasets) aged 18-94 years, the primary aim of this study was to offer a comprehensive description of normative age-related white matter trajectories in adulthood by comparing relevant curve parameters such as key deflection points and rate of change as well as age prediction accuracy of different dMRI metrics, with a particular focus on relatively novel parameters based on advanced (DKI, NODDI, RSI, SMT mc, and WMTI) and conventional (DTI) diffusion models of white matter coherence and microstructure.

First, we estimated the trajectories of each of the diffusion metrics across the age range. Secondly, we utilised three separate methods to compare the age-sensitivity of the diffusion models: i) we used linear mixed effect (lme) models including age, sex, and timepoint, ii) for each model, we ran fits with and without age terms and compared the fit likelihood values using Wilk’s theorem (Wilks, 1938), iii) we used machine learning to predict age based on the diffusion metrics, and compared the prediction accuracy of the models. Thirdly, we looked at the derivatives of each function of the lme models’ age curve to identify the point of change in trajectory for each diffusion metric. Based on previous work characterising age differences and longitudinal changes with a range of diffusion MRI metrics (Benitez et al., 2018; Falangola et al., 2008; Jelescu et al., 2015; Kodiweera et al., 2016; Reas et al., 2017; Westlye et al., 2010), we expected the included metrics to show curvilinear relationships with age, with varying trajectories and deflection points possibly reflecting differential involvement and rate of change of the putative biological underpinnings during the different phases of brain ageing.

## 2. Methods and material

### 2.1. Description of sample

The initial sample included 754 multi-shell datasets of healthy participants from two integrated studies; the Tematisk Område Psykoser (TOP) (Tønnesen et al., 2018) and StrokeMRI (Richard et al., 2018). Following the removal of 52 datasets after quality checking (QC, see section 2.4), the final sample comprised 702 scans from 573 individuals, including longitudinal data (two time-points with 15.2 months interval) for 129 of the participants. Demographic information is summarised in Table 1 and Figure 1.

**Table 1.** Demographics of descriptive statistics pertaining to the study sample. N refers to datasets.

|  | Age |  |  |
| --- | --- | --- | --- |
| | Mean ± SD | Min | Max |
| **Full (mixed) sample (n = 702)** | 50.86 ± 16.61 | 18.52 | 94.67 |
| Male (301, 42.88%) | 49.45 ± 17.48 | 18.52 | 92.05 |
| Female (401, 57.12%) | 51.92 ± 15.86 | 18.63 | 94.67 |
| **Cross-sectional sample (n = 444)** | 47.61 ± 16.59 | 18.52 | 94.67 |
| Male (214, 48.20%) | 46.75 ± 16.71 | 18.52 | 92.05 |
| Female (230, 51.80%) | 48.57 ± 16.51 | 18.63 | 94.67 |
| **Longitudinal sample (n = 258)** | 56.60 ± 15.03 | 20.30 | 85.82 |
| Male (44, 35.11%) | 55.72 ± 17.78 | 20.30 | 85.82 |
| Female (85, 65.89%) | 55.65 ± 13.70 | 23.37 | 80.62 |

**Figure 1.**
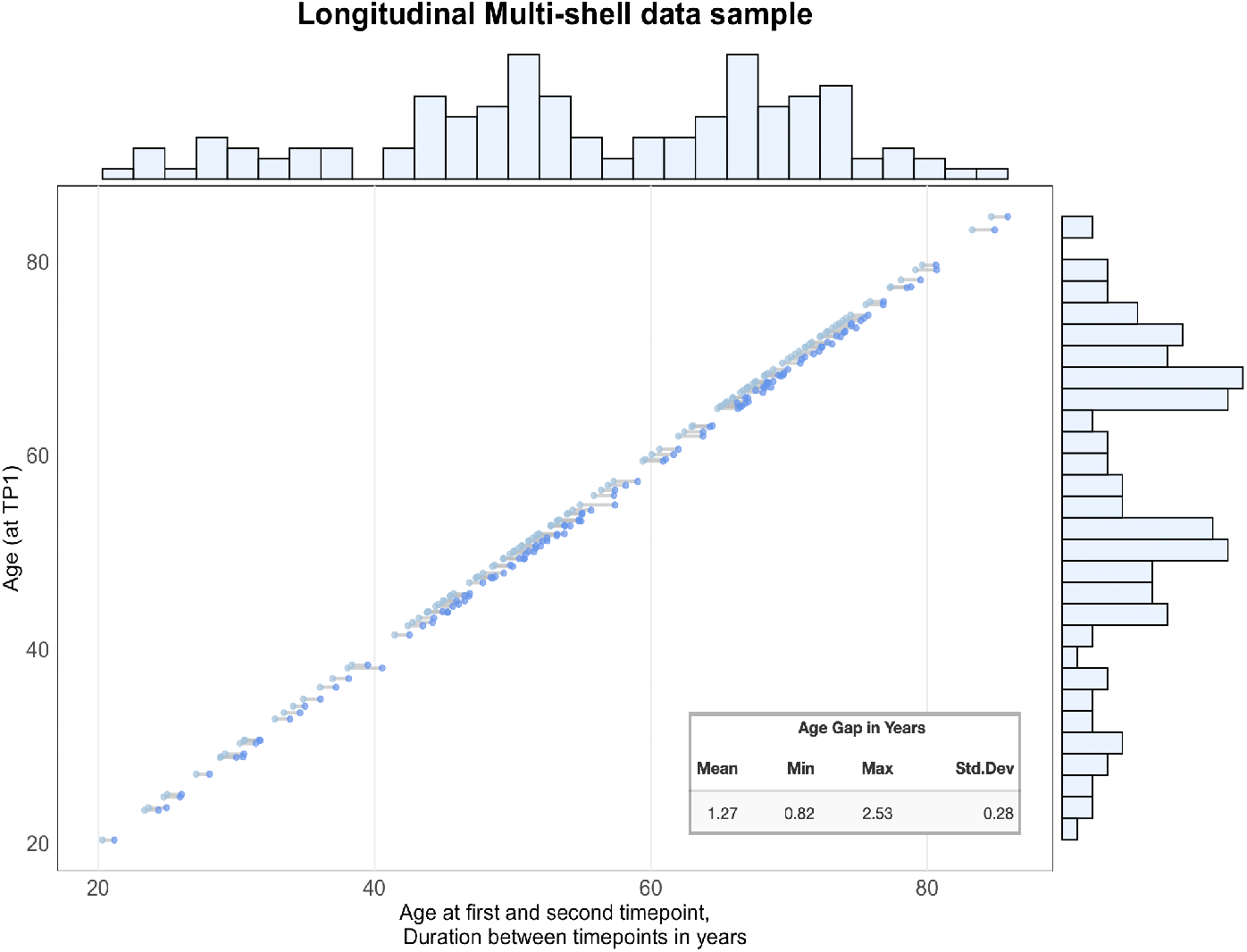
Interval between timepoint 1 and timepoint 2 for complete longitudinal sample, n = 258 (129 subjects). Histogram representing density of data points.

Exclusion criteria included neurological and mental disorders, and previous head trauma. Ethical guidelines followed those in line with the Declaration of Helsinki. The study has been approved by the Regional Ethics Committee and all participants provided written informed consent.

### 2.2. MRI acquisition

Imaging was performed at Oslo University Hospital on a General Electric Discovery MR750 3T scanner with a 32-channel head coil. dMRI data were acquired with a spin echo planar imaging (EPI) sequence with the following parameters: TR/TE/flip angle: 8,150 ms/83.1 ms/90°, FOV: 256 × 256 mm, slice thickness: 2 mm, in-plane resolution: 2 mm. We obtained 10 volumes of *b*=*0* and diffusion weighted data along 60 (*b*=1000 s/mm^2^) and 30 (*b*=2000 s/mm^2^) diffusion weighted volumes. In addition, 7 *b*=0 volumes with reversed phase-encoding direction were acquired for correction of susceptibility distortions.

### 2.3. Diffusion MRI processing

Processing steps followed a previously described pipeline (Maximov et al., 2019), including noise correction (Veraart et al., 2016), Gibbs ringing correction (Kellner et al., 2016), corrections for susceptibility induced distortions, head movements and eddy current induced distortions using topup (http://fsl.fmrib.ox.ac.uk/fsl/fslwiki/topup) and eddy (http://fsl.fmrib.ox.ac.uk/fsl/fslwiki/eddy) (Andersson & Sotiropoulos, 2016). Isotropic smoothing was carried out with a Gaussian kernel of 1 mm^3^ implemented in the FSL function *fslmaths*. DTI was estimated using FSL tool *dtifit* and excluded the b=2000 shell from the fit. Employing the multi-shell data, DKI and WMTI metrics were estimated using Matlab code (https://github.com/NYU-DiffusionMRI/DESIGNER), (Fieremans et al., 2011). NODDI metrics were derived using AMICO in Matlab (https://github.com/daducci/AMICO). SMT mc metrics were estimated with the original code (https://github.com/ekaden/smt). RSI metrics were estimated using in-house Matlab tools.

We selected 20 scalar metrics from the six models (DTI, DKI, NODDI, RSI, SMT mc, WMTI) based on recent studies (Benitez et al., 2018; De Santis et al., 2011; Hope et al., 2019; Jelescu et al., 2015; Kaden, Kelm, et al., 2016; Maximov et al., 2019; Pines et al., 2020). Models were also selected based on feasibility in relation to our acquisition protocol and availability of open source scripts. Figure 2 shows each of the included metrics for one participant, for illustrative purposes. All metrics and their corresponding abbreviations are summarised in Supplementary table 1). Brain age prediction was performed for each model, using all available metrics extracted from a range of regions-of-interest (see section 2.5).

**Figure 2.**
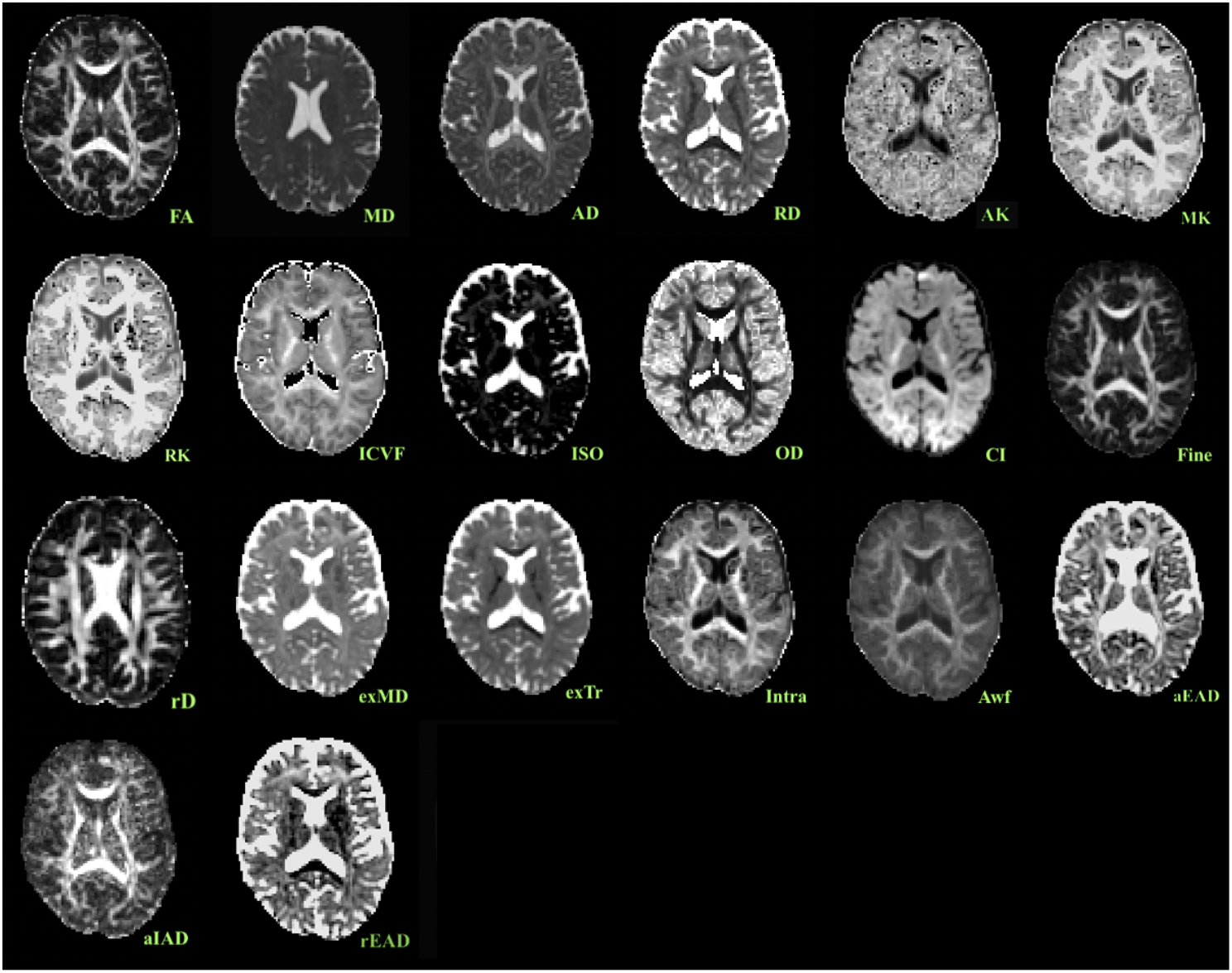
Diffusion metrics from one participant. **DTI**: *FA* (fractional anisotropy), *MD* (mean diffusivity), *AD* (axial diffusivity), *RD* (radial diffusivity). **DKI**: *AK* (axial kurtosis), *MK* (mean kurtosis), *RK* (radial kurtosis). **NODDI**: *ICVF* (intracellular volume fraction), *ISOVF* (isotropic volume fraction), *OD* (oriental dispersion). **RSI**: *CI* (cellular index), *Fine* (FA fine scale/slow compartment), *rD* (restricted diffusivity coefficient). **SMT mc**: *exMD* (extra cellular space), *exTr* (extra-cellular space transverse), *Intra* (intra axonal diffusivity). **WMTI**: *Awf* (axonal water fraction), *aEAD, aIAD* (axial extra and intra axonal diffusivity), *rEAD* (radial extra axonal diffusivity).

### 2.4. Quality checking procedure

We implemented a rigorous QC procedure to ensure data quality was not contaminated by motion, noise, or artefacts. Using a published approach (Roalf et al., 2016), we derived various quality assurance (QA) metrics (see Supplementary material; SI table 2), including temporal-signal-to-noise-ratio (TSNR). Outliers were manually checked and removed if deemed to have unsatisfactory data quality. A total of 52 datasets were removed, leaving the dataset at n = 702 scans. This dataset was put through the same visual inspection. As an additional step, images were manually inspected if TSNR Z scores deviated minus or plus 2.5 standard deviations from the mean. Following this step, the final dataset remained at 702 scans from 573 individuals.

**Table 2.**
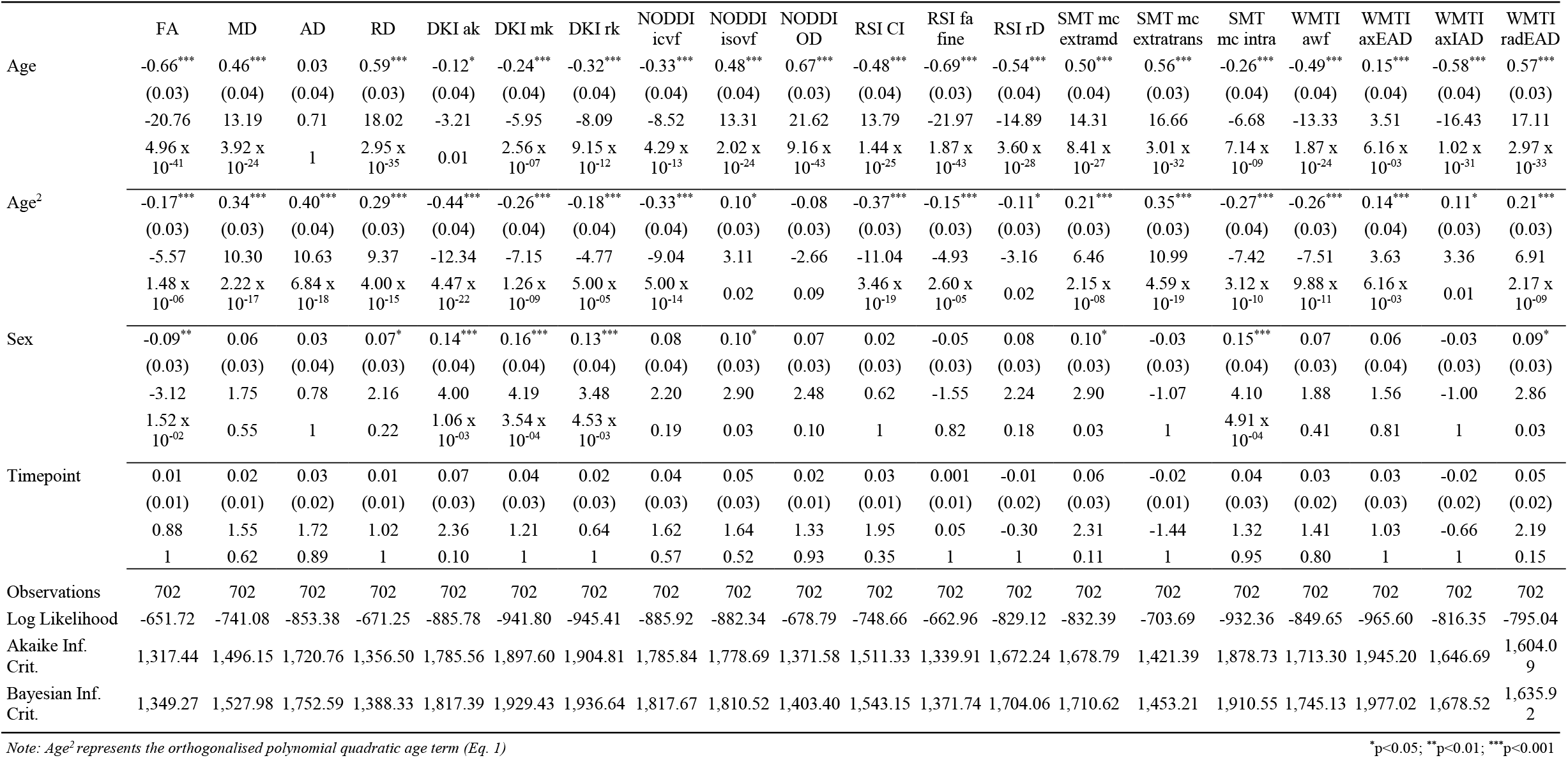
**Linear mixed effect model results for each metric, where variables are** *displayed with corresponding fixed effect estimates (β), (standard error), t-statistic, and FDR corrected P value*.

### 2.5. Tract-Based-Spatial-Statistics

Voxelwise statistical analysis of the FA data was carried out using Tract-Based Spatial Statistics (TBSS) (S. M. Smith et al., 2006), as part of FSL (S. M. Smith et al., 2004). First, FA images were brain-extracted using BET (S. M. Smith, 2002) and aligned into a common space (FMRI58_FA template) using the nonlinear registration tool FNIRT (Andersson, Jenkinson, & Smith., 2007; Jenkinson et al., 2012), which uses a b-spline representation of the registration warp field (Rueckert et al., 1999). Next, the mean FA image of all subjects was created and thinned to create a mean FA skeleton that represents the centres of all tracts common to the group. Each subject’s aligned FA data was then projected onto this skeleton. The mean FA skeleton was thresholded at FA > 0.2. This procedure was repeated for all metrics. *fslmeants* was used to extract the mean skeleton and 20 regions of interest (ROI) based on a probabilistic white matter atlas (JHU) (Hua et al., 2008) for each metric. Including the mean skeleton values, 420 features per individual were derived (20 metrics x 20+1 ROIs). Of these, 20 metrics were used for fitting of age curve trajectories, lme analysis, and Wilk’s theorem analysis, while all 420 MRI features were used for age prediction. Number of MRI features can be found in Table 4. Additional voxelwise analysis were carried out on the 573 participants (excluding longitudinal measures) using the FSL tool Randomise with permutation-based statistics (Winkler et al., 2014) and threshold-free cluster enhancement method (TFCE; (S. Smith & Nichols, 2009)). 5000 permutations were run, where each diffusion metric was tested for its association with age. TBSS fill was used to create voxelwise statistical maps for each metric, which can be found in SI Figure 10.

### 2.6. Diffusion metric reproducibility

The validity and sensitivity of the different diffusion models essentially rely on the richness, quality and specific properties of the data used for model fitting. In order to assess the reproducibility of the included advanced metrics (Maximov et al., 2015), we estimated the dMRI models using data obtained from different acquisition schemes varying the number of directions and maximum *b* value in 23 healthy participants with mean age 35.77 years (SD = 8.37, 56.5% women). This represented a sub-sample of the full sample. The following three acquisition schemes were compared: i) b=[1000,2000] with [60,30] directions, which is identical to the acquisition scheme used in the main analysis, ii) b=[1000,2000] with [60,60] directions and iii) b=[1000,2000,3000] with [60,60,60] directions. For each scheme we processed the data using an identical pipeline (Maximov et al., 2019) as described above and extracted the mean skeleton values for each metric. The comparisons between acquisition protocols were performed using box plots (SI Figure 4), scatterplots with age as a function of mean skeleton values (SI Figures 5), and Pearson’s correlation coefficient plots, where protocol 1 is factored by protocol 3 (SI Figures 6).

**Figure 3.**
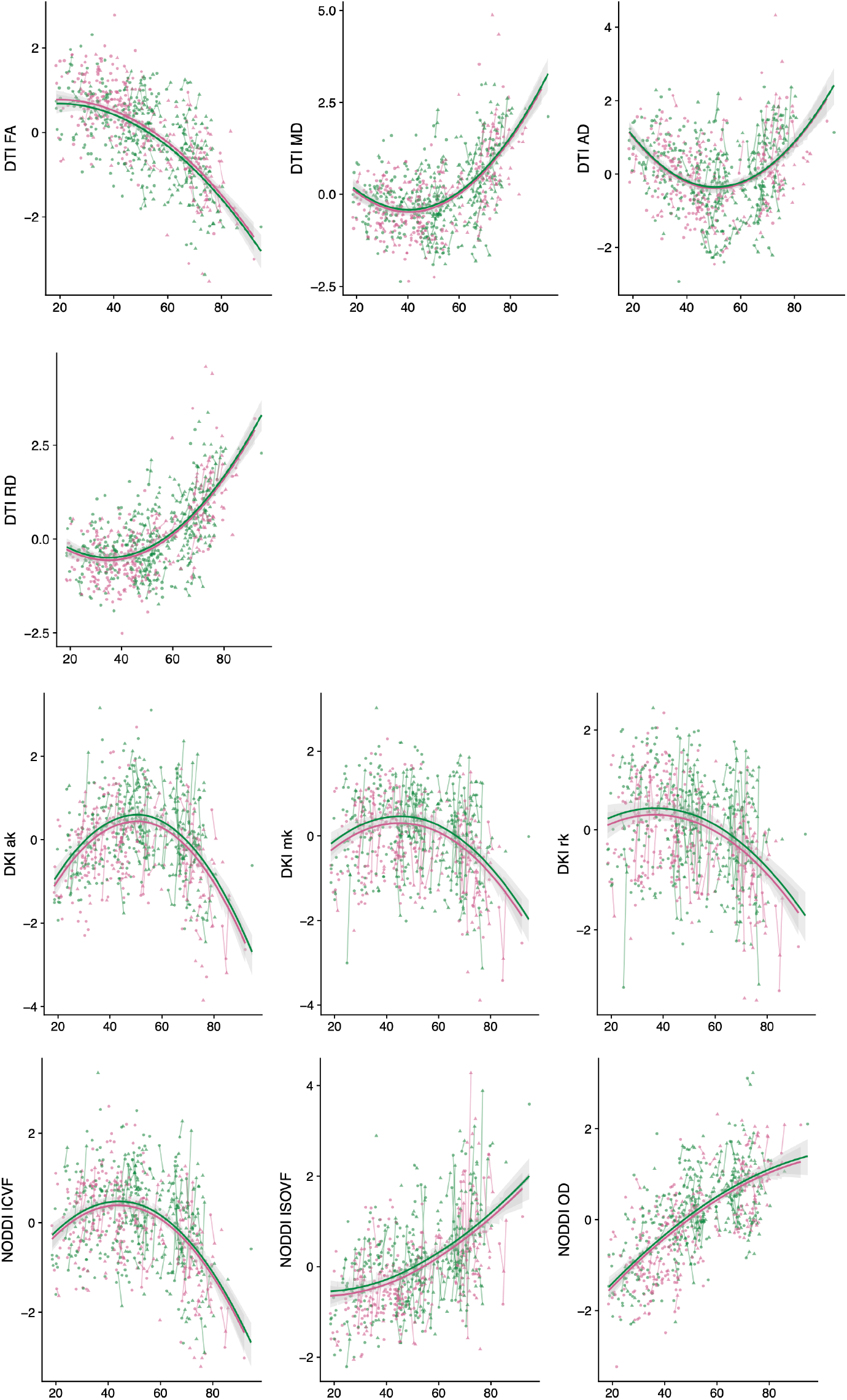
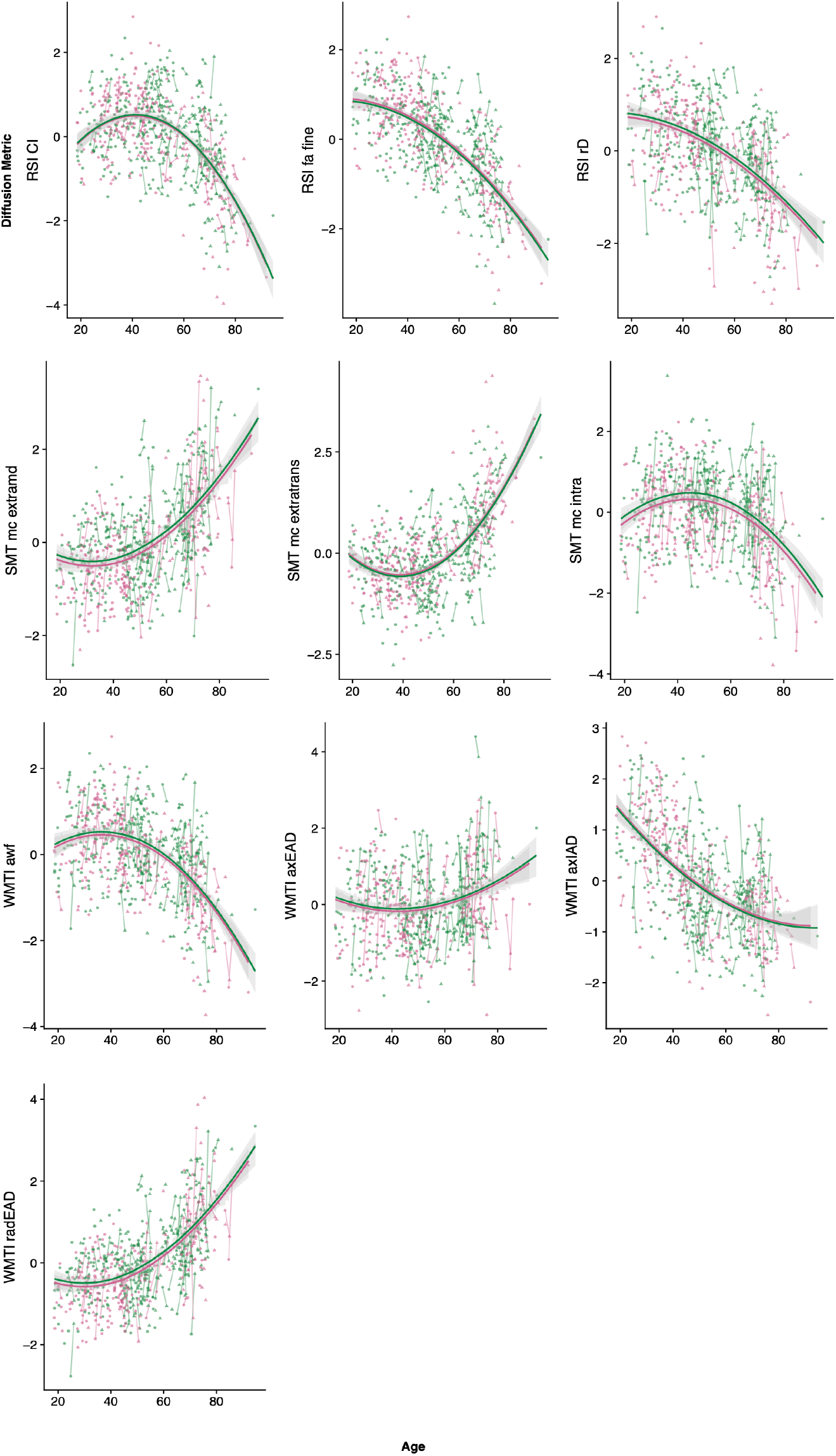
Age curves where each diffusion metric’s standardised (z-score) mean skeleton value (y-axis) is plotted as a function of age (x-axis). Fitted lines made with lme-derived predicted values. Shaded areas represent 95% CI. Points connected by lines represent longitudinal data where circle is TP1 and triangle is TP2. Male subjects are represented by pink and female subjects by green.

**Figure 4.**
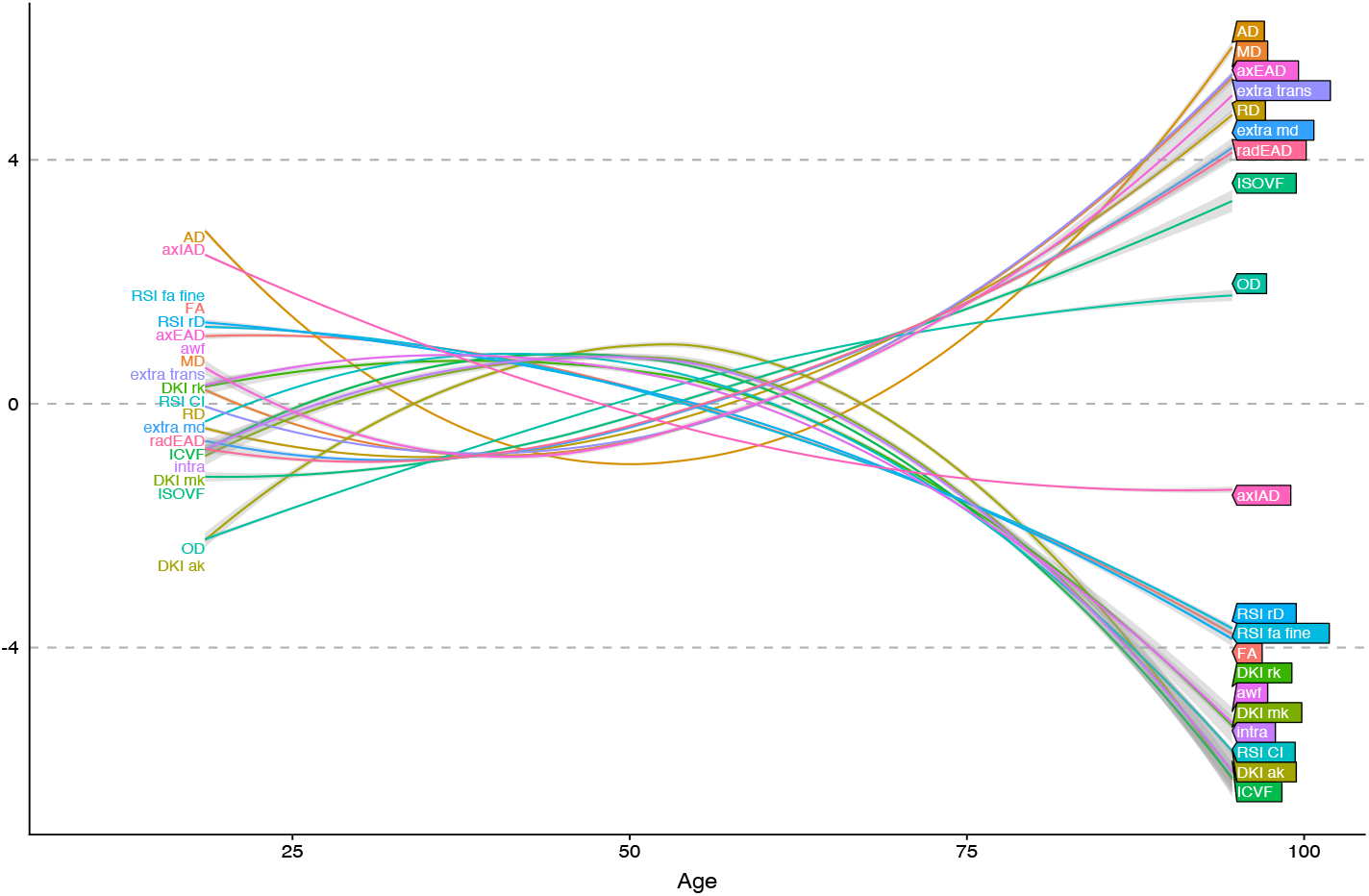
Plot displaying all lme-model derived age curves from Figure 3 in standardised form.

**Figure 5.**
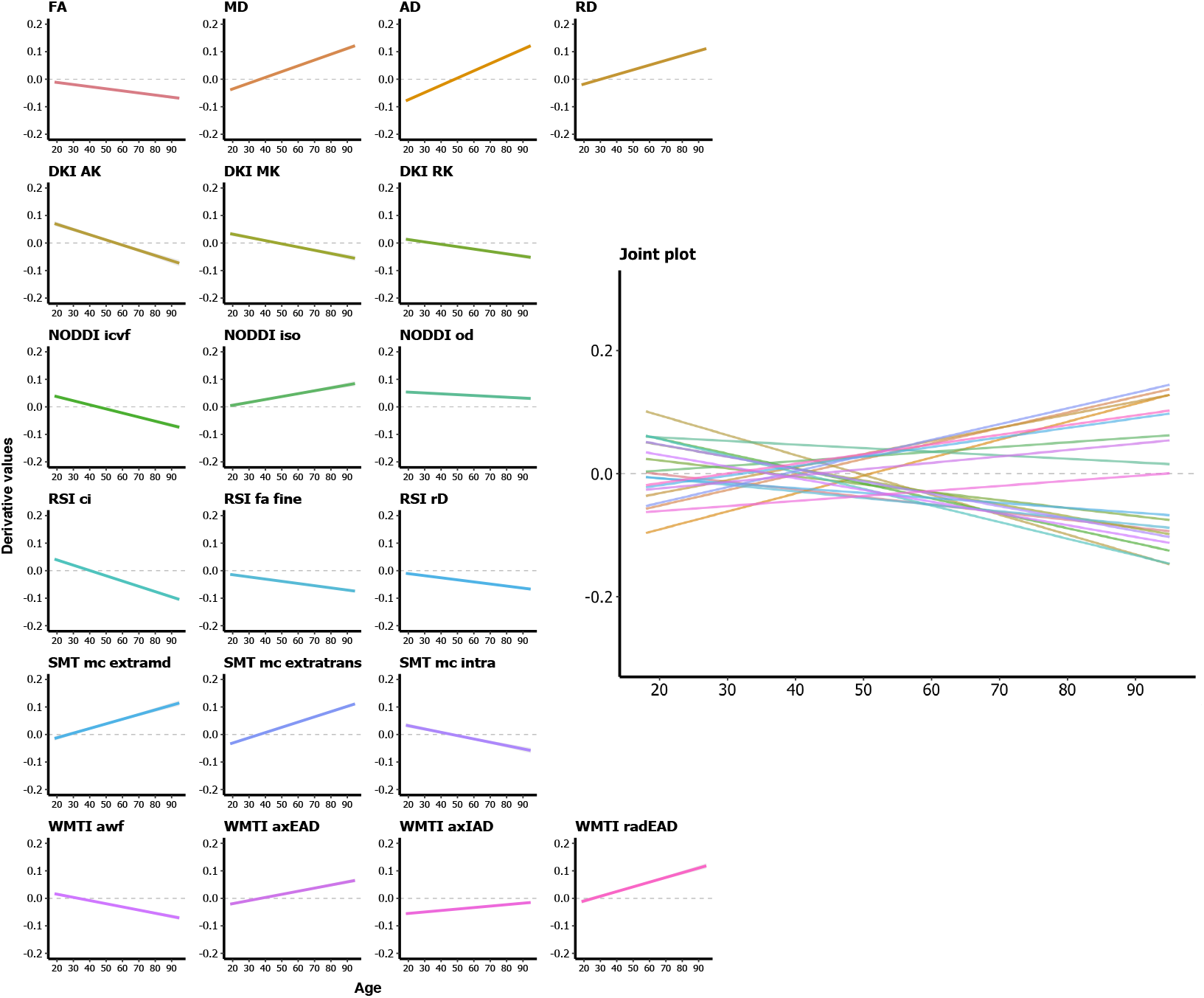
The derivative for each diffusion model, providing the estimated rate of change at every point. The point on the x-axis where the fitted line crosses 0 on the y-axis represents the turning point of the age trajectory for each metric.

**Figure 6.**
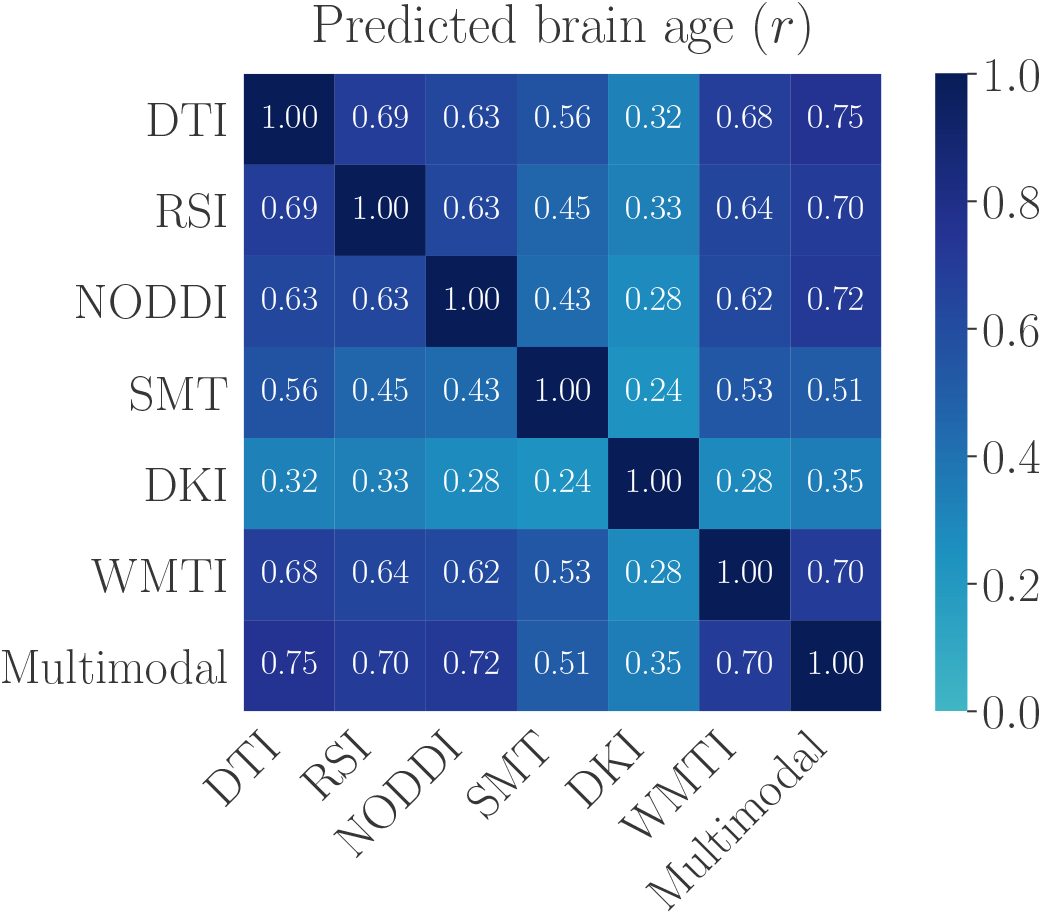
Correlation matrix for predicted brain age of each modality and the multimodal model. To account for age-bias (Le et al., 2018; S. M. Smith, Vidaurre, et al., 2019), the predicted age values were residualised for chronological age using linear models.

### 2.7. Statistical analysis

All statistical analyses were carried out using the statistical environment R, version 3.6.0 (www.r-project.org/) (R Core Team, 2012) and Python 3.7.0 (www.python.org/).

### 2.8. Linear mixed effects models (lme)

To investigate the relationship between age and global mean skeleton values for each diffusion metric, lme analyses were performed using the *lme* function (Bates & Pinheiro, 1998) in R (R Core Team, 2012). In fitting the model, we scaled (z normalised) each variable and entered age, orthogonalised age^2^, sex, and timepoint (TP) as fixed effects. Subject ID was entered as a random effect. For each diffusion metric M, we employed the following function:

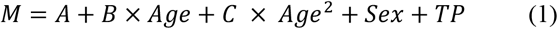

where A is the intercept, B is the age coefficient, and C is the coefficient of the orthogonalised quadratic age term (expressed as *poly(age,2)[,2]* in R). Age curves were obtained with predictions from the fitted model using the *predict* function in R and used for age curve trajectory figures. Visual inspection of residual plots did not reveal any obvious deviations from homoscedasticity or normality. The significance threshold was set at *p* < 0.05, and the results were corrected for multiple comparisons using the false discovery rate (FDR) adjustment (Benjamini & Hochberg, 1995).

To investigate the rate of change for each of the age curves at any point, we calculated their derivatives using numerical differentiation with finite differences (Burden & Faires, 2011). To compare the age-sensitivity of the models, we ran lme fits with and without age terms, and calculated the difference in likelihood ratios (Glover & Dixon, 2004). The significance of the age dependence was calculated using Wilk’s theorem (Wilks, 1938) as 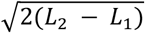, where L_2_ is the likelihood ratio obtained from the models with age terms, and L_1_ is the likelihood ratio obtained from the models without age terms.

### 2.9. Brain-age prediction

The XGBoost regressor model was used to run the brain age prediction (https://xgboost.readthedocs.io/en/latest/python/index.html), including a decision-tree-based ensemble algorithm that has been used in recent large-scale brain age studies (A.-M. G. de Lange et al., 2019; Kaufmann et al., 2019). Parameters were set to max depth = 3, number of estimators = 100, and learning rate = 0.1 (defaults). For each diffusion model (DTI, DKI, NODDI, RSI, SMT mc, WMTI), predicted age was estimated in a 10-fold cross validation, assigning a model-specific brain age estimate to each individual, as well as a multimodal brain age estimate based on all diffusion features. To investigate the prediction accuracy of each model, correlation analyses were run for predicted versus chronological age, and model-specific R^2^, root mean square error (RMSE) and mean absolute error (MAE) were calculated. To statistically compare the prediction accuracy of the models, Z tests for correlated samples (Zimmerman, 2012) were run on the model-specific correlations between predicted and chronological age in a pairwise manner using

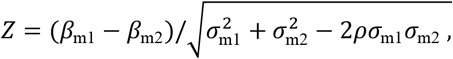

where “m1” and “m2” represent model 1 and model 2, the β terms represent the beta value from the regression fit, the σ terms represent their errors, and ρ represents the correlation between the two sets of associations. In order to assess the complementary value of the different models, we computed the correlations between the brain age predictions (Figure 6). The predictions were first corrected for age-bias using linear models (Le et al., 2018), and the residuals were used in the correlation analysis.

To evaluate the importance of each diffusion modality in the multimodal model, we ran an additional prediction model including only mean-skeleton values to reduce the number of highly correlated features in the regressor input, and calculated a) the proportion of the total *weight* contributed by each modality, where weight represents the number of times a feature is used to split the data across all trees, and b) *gain* values, which represent the improvement in accuracy added by a feature to the branches it is on. To assess the significance of the general model performance, average RMSE was calculated for the multimodal model using cross validation with ten splits and ten repetitions and compared to a null distribution calculated from 1000 permutations.

## 3. Results

### 3.1. Diffusion metric reproducibility

The reproducibility of the estimated diffusion metrics based on data obtained with different acquisition schemes (described in 2.6) revealed overall high correlations between the mean skeleton values for all the model metrics. Highest overall reproducibility was observed for NODDI OD (*r*(22) = 0.96, *p* < 0.001) and RSI rD (*r*(22) = 0.97, *p* < 0.001). The lowest reproducibility was observed for WMTI radEAD (*r*(22) = 0.44, *p* = 0.597). Supplementary Table 4 and Supplementary Figures 4, 5, 6, and 7 show the results from the comparisons.

**Figure 7.**
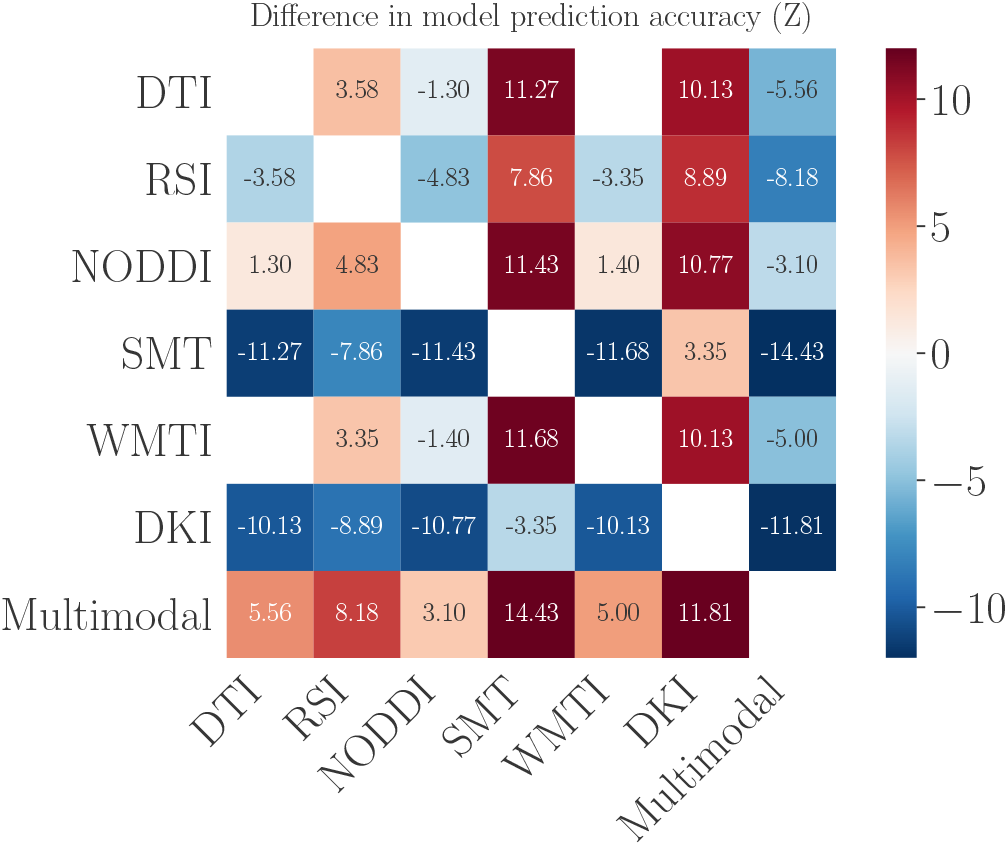
Matrix showing pairwise differences between the model prediction accuracies (correlations between predicted and chronological age), based on z tests for correlated samples.

### 3.2. Age trajectories

Figure 3 shows the linear mixed effect model-derived age curves for each diffusion metric plotted as a function of age, where age curves are fitted with the predicted values of the lme models. Figure 4 shows all lme model-derived age curves from Figure 3 in standardised form in one plot. Figure 5 shows the derivatives of the lme fits, providing the estimated rate of change at every point (of age), including the point of change in trajectory for each model curve and the steepness of the turning point. Correlations between the metrics are available in the supplementary material (SI Figures 2 and 3) for both raw and standardised values respectively.

### 3.3. Comparing age curves

Figure 3 shows the estimated age curves for all metrics. Briefly, FA decreased steadily after the age of 30, with a steeper decline after the age of 50. MD, AD, and RD followed a reverse profile, with decreases in diffusivity until the 40’s, whereby the trajectories subsequently increased thereafter. DKI metrics revealed curvilinear trajectories, with NODDI ICVF, RSI CI, SMT mc intra, and WMTI awf metrics following similar trajectories. RSI rD, NODDI ISOFV, RSI FA fine, and WMTI axIAD metrics followed decreasing trajectories from the offset. SMT mc extramd and extratrans, and WMTI radEAD followed similar trajectories to MD and RD. NODDI OD revealed a steady increase until older age where the slope stabilised thereafter. Lastly, WMTI axEAD showed u-trajectories.

### 3.4. Age sensitivity estimated using lme models

Results from the lme models revealed significant main effects of age on the global mean skeleton values for all diffusion metrics (see Table 2). An examination of the fixed effects estimates (*β*) and t-statistics for the age term allows for interpretation of the extent and direction of the linear association with age. Overall, the FA fine compartment of the RSI model was most sensitive to age (*β*(125) = −0.69, t = −21.97, *p* < 0.001). NODDI OD was the second most sensitive to age (*β*(125) = 0.67, t = 21.62, *p* < 0.001). The model least sensitive to age was DTI AD (*β*(125) = 0.03, t = 0.71, *p* = 1). For conventional DTI metrics, FA was the most age sensitive (*β*(125) = −0.66, t = −20.76, *p* < 0.001). No main effects of timepoint survived correction for multiple comparisons.

### 3.5. Age sensitivity estimated using Wilk’s theorem

Table 3 shows the strength of the overall age variation for each metric estimated by the difference in likelihood values (described in Section 2.8). All metrics showed significant age dependence, with RSI FA fine as the most age sensitive (z = 18.79), followed by NODDI OD (z = 18.55) and DTI-based FA (z = 18.12). WMTI axEAD (z = 4.65) was the least age-dependant metric.

**Table 3.** Likelihood values from the lme models without age terms (L1) and with age terms (L2). The significance of the age dependence is estimated by the difference in likelihood values using Wilk’s theorem. FDR corrected p-values = *p*^corr^.

| Model | | $L_1$ | $L_2$ | Difference (z) | p-value | $p^{\text{corr}}$ |
| --- | --- | --- | --- | --- | --- | --- |
| DTI | FA | -815.86 | -651.72 | 18.12 | $5.22 \times 10^{-72}$ | $1.04 \times 10^{-70}$ |
| | MD | -848.36 | -741.08 | 14.65 | $2.55 \times 10^{-47}$ | $5.10 \times 10^{-46}$ |
| | AD | -900.66 | -853.38 | 9.72 | $2.93 \times 10^{-21}$ | $5.86 \times 10^{-20}$ |
| | RD | -820.44 | -671.25 | 17.27 | $1.62 \times 10^{-65}$ | $3.24 \times 10^{-64}$ |
| DKI | AK | -952.44 | -885.78 | 11.55 | $1.12 \times 10^{-29}$ | $2.25 \times 10^{-28}$ |
| | MK | -977.09 | -941.80 | 8.40 | $4.71 \times 10^{-16}$ | $9.42 \times 10^{-15}$ |
| | RK | -981.65 | -945.41 | 8.51 | $1.83 \times 10^{-16}$ | $3.65 \times 10^{-15}$ |
| NODDI | ICVF | -948.54 | -885.92 | 11.19 | $6.40 \times 10^{-28}$ | $1.28 \times 10^{-26}$ |
| | ISOVF | -957.61 | -882.34 | 12.27 | $2.06 \times 10^{-33}$ | $4.13 \times 10^{-32}$ |
| | OD | -850.84 | -678.79 | 18.55 | $1.90 \times 10^{-75}$ | $3.80 \times 10^{-74}$ |
| RSI | CI | -866.73 | -748.66 | 15.37 | $5.28 \times 10^{-52}$ | $1.06 \times 10^{-50}$ |
| | FA fine | -839.53 | -662.96 | 18.79 | $2.07 \times 10^{-77}$ | $4.15 \times 10^{-76}$ |
| | rD | -922.24 | -829.12 | 13.65 | $3.62 \times 10^{-41}$ | $7.24 \times 10^{-40}$ |
| SMT mc | Extra md | -929.01 | -832.39 | 13.90 | $1.10 \times 10^{-42}$ | $2.20 \times 10^{-41}$ |
| | Extra trans | -848.79 | -703.69 | 17.03 | $9.71 \times 10^{-64}$ | $1.94 \times 10^{-62}$ |
| | Intra | -973.21 | -932.36 | 9.04 | $1.82 \times 10^{-18}$ | $3.64 \times 10^{-17}$ |
| WMTI | AWF | -942.66 | -846.37 | 13.88 | $1.52 \times 10^{-43}$ | $3.04 \times 10^{-41}$ |
| | axEAD | -973.15 | -962.32 | 4.65 | $1.98 \times 10^{-05}$ | $3.97 \times 10^{-04}$ |
| | axIAD | -930.26 | -816.35 | 15.09 | $3.38 \times 10^{-50}$ | $6.76 \times 10^{-49}$ |
| | radEAD | -922.92 | -795.04 | 15.99 | $2.91 \times 10^{-56}$ | $5.81 \times 10^{-55}$ |

**Table 4.** Number of MRI variables (corresponding to the sum of metric features), root mean square error (RMSE), mean absolute error (MAE), correlation between predicted and chronological age (Pearson’s r), and R^2^ for each of the models. CI = confidence interval.

| Model | MRI variables | RMSE | MAE | $r$ [95% CI] | $R^2$ [95% CI] |
| --- | --- | --- | --- | --- | --- |
| DTI | 84 | 9.35 | 7.30 | 0.83 [0.80, 0.85] | 0.68 [0.64, 0.72] |
| DKI | 63 | 12.19 | 9.82 | 0.68 [0.64, 0.72] | 0.46 [0.41, 0.52] |
| NODDI | 63 | 9.15 | 7.31 | 0.83 [0.81, 0.86] | 0.70 [0.65, 0.74] |
| RSI | 63 | 9.84 | 7.68 | 0.81 [0.78, 0.83] | 0.65 [0.61, 0.69] |
| SMT mc | 63 | 11.30 | 9.01 | 0.73 [0.70, 0.76] | 0.54 [0.50, 0.58] |
| WMTI | 84 | 9.37 | 7.40 | 0.83 [0.80, 0.85] | 0.68 [0.64, 0.72] |
| Multimodal | 420 | 8.80 | 6.99 | 0.85 [0.83, 0.87] | 0.72 [0.69, 0.76] |

### 3.6. Age sensitivity estimated using brain age

The model performances for the multimodal and model-specific brain age predictions are shown in Table 4. SI Figures 8 and 9 show the associations between predicted age and chronological age for each of the models. Figure 6 shows the pairwise correlations between predicted age for each model. Pairwise differences in the age prediction accuracy of the models are shown in Figures 7 and 8. SI Figure 1 shows the RMSE of the multimodal model prediction compared to a null distribution obtained from calculating 1000 permutations.

**Figure 8.**
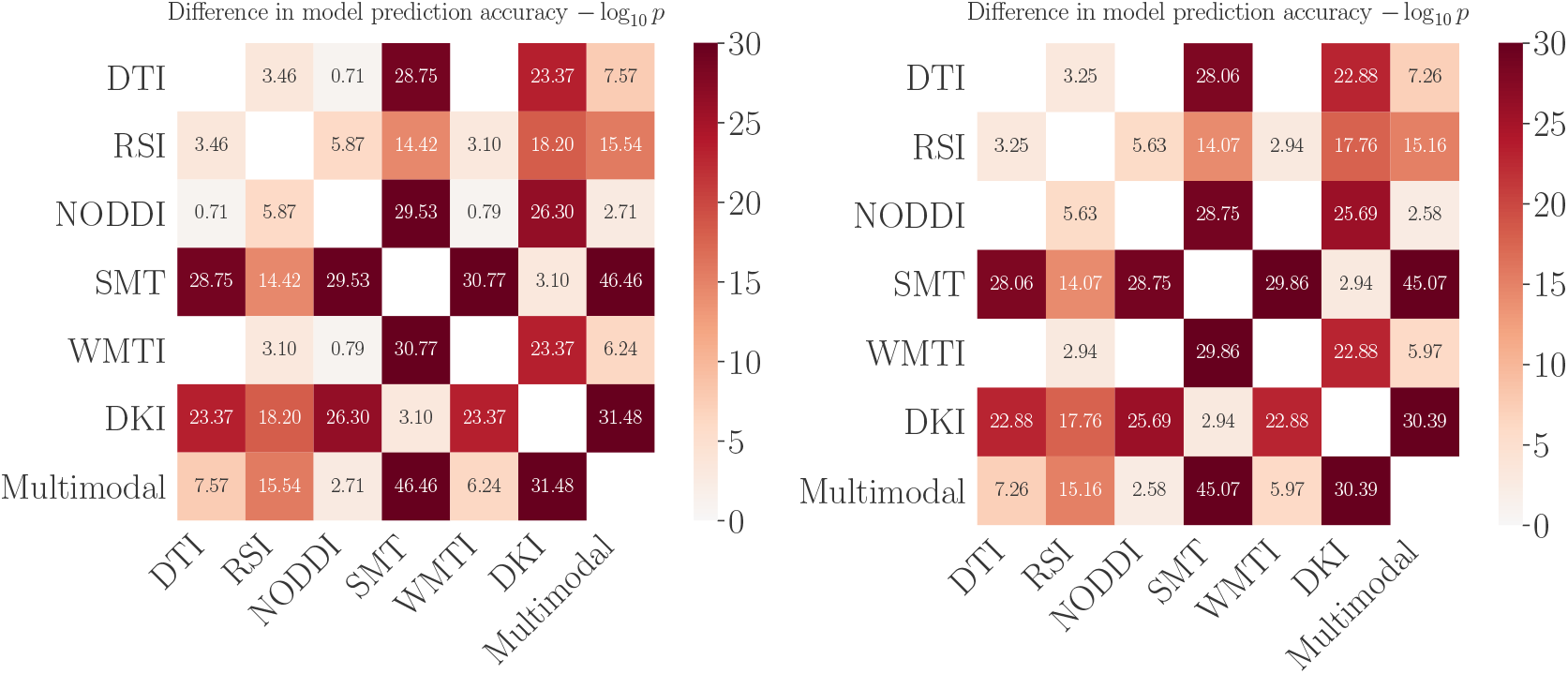
Log10(*p*) values of the pairwise differences between the model prediction accuracies. Higher numbers represent more significant differences. Left: uncorrected *p*-values. Right: *P*-values corrected for multiple comparisons using FDR, with non-significant (> 0.05) values masked out.

As visible from Table 4, the multimodal model showed the most accurate age prediction (*r* = 0.85, *p* < 0.001, 95% CI = [0.83, 0.87]), while the DKI model performed the worst (*r* = 0.68, *p* < 0.001, 95% CI = [0.64, 0.72]). As shown in Figures 7 and 8, the multimodal prediction accuracy was significantly higher than the accuracy of each of the other models, with the largest difference seen between the multimodal model and DKI. The differences in prediction accuracy between DTI and RSI, and WMTI and NODDI did not survive correction for multiple comparisons. Figure 6 showed correlation coefficients of mean *r* = 0.59 (Std = 0.09) between the DTI, RSI, NODDI, SMT and WMTI predictions, while the DKI showed lower correlations with the other model predictions (mean *r* = 0.29, Std = 0.04).

To evaluate the relative importance of each modality, we ran an additional multimodal model including only mean-skeleton values to reduce the number of highly correlated features in the regressor input. Table 5 shows the total gain and the proportion of weight contributed by each modality to the total weight, indicating their relative contribution in the model training. The results revealed that the machine favoured the NODDI model in the training.

**Table 5.** Feature importance evaluated using a reduced multimodal model that included only mean skeleton values for each modality. Number of MRI variables (corresponding to the sum of metric features), percentage contribution to the total weight, and total gain for each modality.

| Model | MRI variables | % of total weight | Total gain |
| --- | --- | --- | --- |
| DTI | 4 | 20.09 | 163473.25 |
| DKI | 3 | 5.13 | 41747.63 |
| NODDI | 3 | 45.48 | 370129.31 |
| RSI | 3 | 4.85 | 39463.11 |
| SMT mc | 3 | 11.74 | 95534.98 |
| WMTI | 4 | 12.72 | 103545.15 |

## 4. Discussion

Ageing confers a range of structural brain alterations, affecting micro- and macrostructural properties of the neurocircuitry supporting cognitive and other complex brain functions. In the current mixed cross-sectional and longitudinal study, we compared age sensitivity and brain white matter age trajectories across the adult lifespan based on advanced and conventional dMRI models. The results from our comprehensive analysis approach, including age-curve trajectories, linear mixed effects models, Wilk’s theorem analysis, and brain age prediction, showed high age sensitivity for all diffusion metrics, with comparable sensitivity between the highest performing advanced dMRI models and conventional DTI, and a moderate benefit of including all metrics in the same model. The mixed effects analyses and corresponding derivatives revealed variations in age trajectories between models, indicating that they may be sensitive to different underlying aspects of white matter ageing.

Our results showed that FA plateaued around the third decade with a steady decline in slope following the age of ∼40, and an accelerated decrease in senescence (Figure 3). The other DTI metrics of MD, AD, and RD revealed decreases in diffusivity up until the 40-50-year age mark, where the trajectories subsequently increase following a steady period. While these results to a large extent correspond with trajectories observed in previous studies (Cox et al., 2016; Davis et al., 2009; Westlye et al., 2010), a more defined inverted U-shape (Westlye et al., 2010) was less prominent in our study, likely due to a lack of younger participants below the age of 20. Interestingly, FA based on the relatively simple DTI model utilising only single-shell data offered one of the highest sensitivities to age, supporting that DTI provides sensitive measures of gross white matter anatomy and neuropathological changes (A. L. Alexander et al., 2008). The characteristic curvilinear trajectories of lifespan differences in conventional DTI metrics (Westlye et al., 2010) have previously been suggested to reflect a combination of protracted myelin-related maturation during childhood, adolescence and early adulthood (Lebel et al., 2008; Tamnes et al., 2010) and subsequent myelin loss during adulthood and senescence (Bartzokis et al., 2004). However, DTI metrics are unable to differentiate between intra-and extra-axonal compartments, and, in addition to the idiosyncratic changes in myeloarchitecture, they may be influenced by individual differences and changes in gross fiber architecture (e.g. crossing fibres) and axonal packing and density (Paus, 2010; Simmonds et al., 2014). The specific biological interpretation of DTI metrics essentially depends upon the local fiber architecture, and signal changes from DTI require careful interpretation, as the exact neurobiological underpinnings cannot be directly inferred. While speculative, utilising advanced dMRI models in addition to conventional DTI may provide more specificity in the interpretation of the results, and improve the descriptive precision of the tissue pathology by disentangling the various biological sources that are happening concurrently.

While several of the advanced dMRI models showed comparable results to DTI in terms of age sensitivity, they also showed visibly different age trajectories (Figure 3), including variation in turning points (Figure 4), indicating the age at which anisotropy and diffusivity measures change direction, and gradient of change (Figure 5), indicating rate of decline. The variation in turning points and gradient of change calculated using the derivates of each model informs us about the estimated rate of change at specific ages, in addition to the differential sensitivity between different metrics during different life phases. Although diffusion imaging cannot give direct access to neuronal processes on a cellular level, the varying estimated trajectories in advanced dMRI models potentially reflect differential involvement of the putative biological underpinnings during the different phases of brain ageing. Thus, metric-specific differences may reflect age-related pathological changes behind each dMRI model, helping us better pinpoint the age at which decline in white matter microstructure begins, which has important implications for interventive strategies aimed at promoting healthy ageing.

Although recent research has validated FA and RD metrics of DTI as being sensitive markers to myelin (Lazari & Lipp, 2020), caution must be exerted in interpreting specific underlying biology on the basis of DTI alone (Novikov et al., 2018). With this in mind, combining tissue models such as NODDI, WMTI, RSI, and SMT mc may hold promise in jointly reflecting measures more relatable to the neurobiological underpinnings of brain ageing. The WMTI metrics for example have been validated for reflecting underlying biology both *in vivo* (Jelescu et al., 2015, 2016) and *ex vivo* (Falangola et al., 2014; Kelm et al., 2016). WMTI awf was found to relate to axonal density, whereas WMTI radEAD to some extent describes the degree of myelination (Kelm et al., 2016) and relates to the extracellular environment filled with interstitial fluid and circulating macromolecules, as well as blood vessels and perivascular spaces (Nicholson & Hrabětová, 2017). The parameter maps from the NODDI model have been claimed to exhibit a spatial pattern of tissue distribution consistent with the known brain anatomy (Zhang et al., 2012), with existing maps showing the expected pattern of neurite density (Jespersen et al., 2010), serving as an example of the feasibility provided by advanced diffusion models to disentangle neurite density and orientation dispersion, two major factors contributing to FA (Zhang et al., 2012). The RSI model diameter calculations have been shown to correspond with the diameter of unmyelinated and myelinated axons in the rat brain (White et al., 2013), suggesting a direct biological interpretation. Likewise, histological analyses have shown that the SMT mc microscopic diffusion indices offer direct sensitivity to pathological tissue alterations (Kaden et al. 2016). While not a tissue model, DKI provides a specific measure of cellular compartments and membranes and is relatively unconfounded by concentration of macromolecules, potentially providing a more specific indicator of tissue properties than conventional DTI (Jensen et al., 2005).

In theory, the partly non-overlapping assumptions and biophysical properties of the different diffusion MRI models offer a more comprehensive and complete view of the manifold biological processes in brain development, ageing, and disorders when considered jointly. In general, our findings of higher age prediction accuracy when combining different models supports this view. However, not surprisingly, the relatively high correlations and similar age-related trajectories of several of the different metrics also suggest a certain level of redundancy. Further studies are needed to test the hypothesis that combining various diffusion MRI models of brain macro-and microstructure increases the feasibility and precision of multimodal data-driven brain phenotyping approaches (e.g. “fingerprinting”) towards more specific clinical applications and prediction (Alnæs et al., 2018). With this in mind, including the advanced models may not only improves specificity compared to conventional DTI, but potentially provides additional information related to changes in myelination and axonal rewiring, while specifically modelling microstructural features typically conflated by DTI, such as neurite density, axonal diameter, and neurite orientation dispersion (D. C. Alexander et al., 2019). Further research is needed to validate and develop dMRI models to better reflect the different biological and geometrical properties of white matter. If assumptions of underlying microstructure are valid, these advanced models represent a promising contribution to the investigation of brain development and ageing, and aberrant brain biology in various clinical conditions (D. C. Alexander et al., 2019).

While considering a range of diffusion models, it is important to note that each comes with its respective limitations. NODDI has been particularly criticised in recent years, with research suggesting the model assumptions are invalid (Lampinen et al., 2017). NODDI provides estimates of geometric parameters only, with there being an absence of any direct diffusivity estimation (Jelescu et al., 2015). DKI, like DTI, is limited in specificity as it can be affected by different features of tissue microstructure. Thus, the biophysical model that relates DKI parameters directly to white matter microstructure (WMTI, (Fieremans et al., 2011)) was proposed. However, assumptions made in WMTI may be oversimplifying, which could lead to bias in the estimated parameters in addition to reduced information about the microstructure. WMTI parameter estimation accuracy is also said to progressively degrade with higher orientation dispersion (Jelescu et al., 2015).

The SMT mc model overcomes limitations in WMTI (Fieremans et al., 2011) and NODDI (Zhang et al., 2012) as it makes no assumptions about the neurite orientation distribution (Kaden, Kelm, et al., 2016). However, it is limited by assuming that the effective transverse diffusivity inside the neurites is zero, an approximation which may not hold for unmyelinated axons and dendrites (Kaden, Kelm, et al., 2016), due to possible neurite undulation on the microscopic scale (Nilsson et al., 2012). RSI, like most diffusion-based techniques, suffers from low resolution and may best be utilised in supplement to high spatial resolution sequences as part of a multimodal imaging protocol (Brunsing et al., 2017). For example, the DTI model’s limitation of being blind to crossing and bending fibres may be resolved by the RSI model’s multi-direction properties and ability to measure diffusion orientation and length scale (White et al., 2013). Despite the limitations of each model, and possible redundancy between them, assessing age-related white microstructural changes using a combination of diffusion models can be advantageous in order to zero in on idiosyncratic neuroanatomical and microstructural patterns (Alnæs et al., 2018). Biophysical models of WMTI and SMT mc for example, adds the possibility for assessing the separate effect of diffusion in intra-and extra-axonal space (Jelescu & Budde, 2017; Voldsbekk et al., 2020).

Some methodological limitations must also be addressed. One concern is that of averaging over regions of interests and the entire white matter skeleton, which is complicated by the direction and magnitude of age associations varying regionally. Recent findings (Tønnesen et al., 2020) found that the global mean skeleton model outperformed region of interest-based single-metric models, providing evidence for relevant information required for brain age prediction is captured at a global level. Indeed, previous studies have suggested that regional DTI-based indices of brain aging reflect relatively global processes (Penke et al., 2010; Westlye et al., 2010), which is also supported by a genetically informed approach demonstrating that a substantial proportion of the tract-wise heritability is accounted for by a general genetic factor (Gustavson et al., 2019). Secondly, we used FA to generate white matter skeletons. Future research should consider generating white matter skeletons based on advanced diffusion maps that are more resistant to crossing fibres.

Other strengths of the study must also be addressed. TBSS offers robust non-linear registration and skeletonization of individual FA maps, which allows both for subsequent voxel-wise analysis and extraction of ROI based summary stats using a range of white matter atlases. This approach is highly standardized, which promotes reproducibility and future meta-analyses. The direct test of the reproducibility of the included dMRI metrics across different acquisition schemes with a higher number of directions and *b*-values, supported the use of advanced computational dMRI models for data obtained using a clinically feasible acquisition protocol. The combination of advanced dMRI models based on multi-shell data is a key strength, which potentially provides more detailed features of the cellular environment from differential tissue responses elicited by the different *b*-values (Assaf & Basser, 2005; Clark et al., 2002; Pines et al., 2020).

The study also included a relatively large sample and benefitted from all participants having been scanned with the same MRI scanner. Additionally, with cross-sectional studies being limited by between-subject variance and possible cohort effects (Schaie, 2005), the current study profits from a mixed cross-sectional and longitudinal design, where participants can be used as their own baseline (Sexton et al., 2014). However, the longitudinal aspect of our study had some limitations, including the short interval duration, and the low sample size compared to the cross-sectional sample. Consequently, the main results were largely driven by cross sectional data despite the mixed cross-sectional and longitudinal nature of the design. Future research should aim to adopt fully longitudinal designs over several time points in order to evaluate individual differences in change over time, preferably over wide age ranges.

Although the advanced dMRI models offered new insight into age sensitivity (such as the relatively high performance of RSI and NODDI for age prediction) and differences in age trajectories, the biological interpretation of these metrics require further validation. Continued development and validation of more optimal diffusion models that better reflect biological properties of the brain is needed, and future research should take into account the impact of a range of potential factors that may mediate brain and cognitive development (Alnæs et al., 2020) and ageing (Lindenberger, 2014), such as pre- and perinatal events, socio-demographical factors, education, lifestyle, cardiometabolic risk factors, and genetics.

In conclusion, characterising changes in white matter microstructure over the human lifespan is critical for establishing robust models of normative neurodevelopment and ageing, which in turn can help us to better understand deviations from healthy age trajectories. The current study demonstrates that while advanced and conventional dMRI models show comparable age-sensitivity, multi-shell diffusion acquisition and advanced dMRI models can contribute to measuring multiple, complementary aspects of white matter characteristics. Further developing dMRI models in terms of biological tissue specificity remains a challenging yet important goal for understanding white matter development across the human lifespan.

## Acknowledgements

The study is supported by the Research Council of Norway (223273, 249795, 248238, 286838), the South-Eastern Norway Regional Health Authority (2014097, 2015044, 2015073, 2016083, 2018037, 2018076), the Norwegian ExtraFoundation for Health and Rehabilitation (2015/FO5146), KG Jebsen Stiftelsen, ERA-Net Cofund through the ERA PerMed project ‘IMPLEMENT’, and the European Research Council under the European Union’s Horizon 2020 research and Innovation program (ERC StG, Grant 802998).

## Notes

Conflict of interest: none.

https://osf.io/cm8hb/

### Competing Interest Statement

The authors have declared no competing interest.

### Summary of Updates

The revised manuscript has been updated with added diffusion metrics of Axial diffusivity (AD) of the DTI model, and Axonal water fraction (AWF) of the WMTI model. The revised manuscript has also been updated with voxelwise statistical maps. Additionally, WMTI radIAD was removed from the analyses due to the WMTI models inability to derive intra-axonal radial diffusivity, considering that the model assumes zero intra-axonal radial diffusivity.

