## Supplementary material for "White matter microstructure across the adult lifespan: A mixed longitudinal and cross-sectional study using advanced diffusion models and brain-age prediction"

**SI table 1.** Overview of diffusion metrics

| Model | Metrics | Abbreviation used |
|---|---|---|
| **DTI** (Diffusion tensor imaging) | Fractional anisotropy | **FA** |
| | Mean diffusivity | **MD** |
| | Axial diffusivity | **AD** |
| | Radial diffusivity | **RD** |
| **DKI** (Diffusion kurtosis imaging) | Axial kurtosis | **DKI ak** |
| | Mean kurtosis | **DKI mk** |
| | Radial kurtosis | **DKI rk** |
| **NODDI** (Neurite orientation dispersion and density imaging) | Intracellular volume fraction | **NODDI ICVF** |
| | Isotropic volume fraction | **NODDI ISOVF** |
| | Orientation dispersion | **NODDI OD** |
| **RSI** (Restriction spectrum imaging) | Cellular index | **RSI CI** |
| | FA fine scale ("slow") compartment of the small apparent diffusion coefficient of water (ADC) | **RSI fa fine** |
| | Restricted diffusivity coefficient | **RSI rD** |
| **SMT mc** (Spherical mean technique multi-compartment) | Extra-cellular space | **SMT mc extra md** |
| | Extra-cellular space transverse | **SMT mc extra trans** |
| | Intra axonal diffusivity | **SMT mc intra** |
| **WMTI** (White matter tract integrity) | Axonal water fraction | **WMTI awf** |
| | Axial extra axonal diffusivity | **WMTI axEAD** |
| | Radial extra axonal diffusivity | **WMTI radEAD** |
| | Axial intra axonal diffusivity | **WMTI axIAD** |

**SI table 2.** Overview of Quality Assurance (QA) metrics

| QA metric abbreviation | Measure |
|---|---|
| tsnr | Temporal-signal-to-noise-ratio |
| gmean | Global mean intensity |
| drift | Linear drift of signal over time |
| outmax | Outlier measurement maximum |
| outmean | Outlier measurement average |
| meanABSrmsb | Average absolute root-mean square |
| meanRELrmsb | Average relative root-mean square |
| maxABSrmsb | Maximum absolute root-mean square |
| maxRELrmsb | Maximum relative root-mean square |

**SI table 3.** $R^2$m (marginal) showing the variance explained by the fixed effects. $R^2$c (conditional) showing the variance explained by the entire model, including both fixed and random effects.

| Model | $R^2$m | $R^2$c |
|---|---|---|
| DTI FA | 0.46 | 0.97 |
| DTI MD | 0.34 | 0.95 |
| DTI AD | 0.16 | 0.89 |
| DTI RD | 0.43 | 0.97 |
| DKI ak | 0.24 | 0.71 |
| DKI mk | 0.15 | 0.49 |
| DKI rk | 0.14 | 0.45 |
| NODDI ICVF | 0.21 | 0.77 |
| NODDI ISOVF | 0.28 | 0.54 |
| NODDI OD | 0.48 | 0.94 |
| RSI CI | 0.35 | 0.95 |
| RSI fa fine | 0.48 | 0.96 |
| RSI rD | 0.3 | 0.84 |
| SMT mc extramd | 0.34 | 0.73 |
| SMT mc extratrans | 0.41 | 0.96 |
| SMT mc intra | 0.16 | 0.55 |
| WMTI awf | 0.29 | 0.79 |
| WMTI axEAD | 0.05 | 0.54 |
| WMTI axIAD | 0.35 | 0.82 |
| WMTI radEAD | 0.41 | 0.76 |

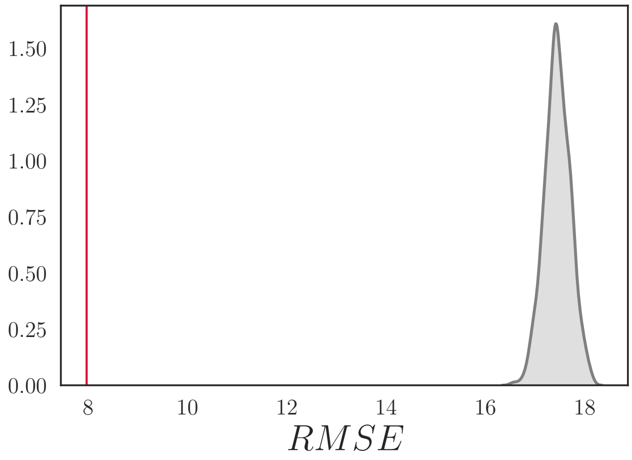


**SI Figure 1.** RMSE distribution. The mean ± SD root mean square error (RMSE) for the multimodal brain age model was 8.30 ± 0.72, based on the cross validation (red vertical line). The null distribution calculated from 1000 permutations is shown in grey, with a mean ± SD of 17.45 ± 0.26. The number of permuted results from the null distribution that exceeded the mean from the cross validation was 0 ($p < 0.001$).

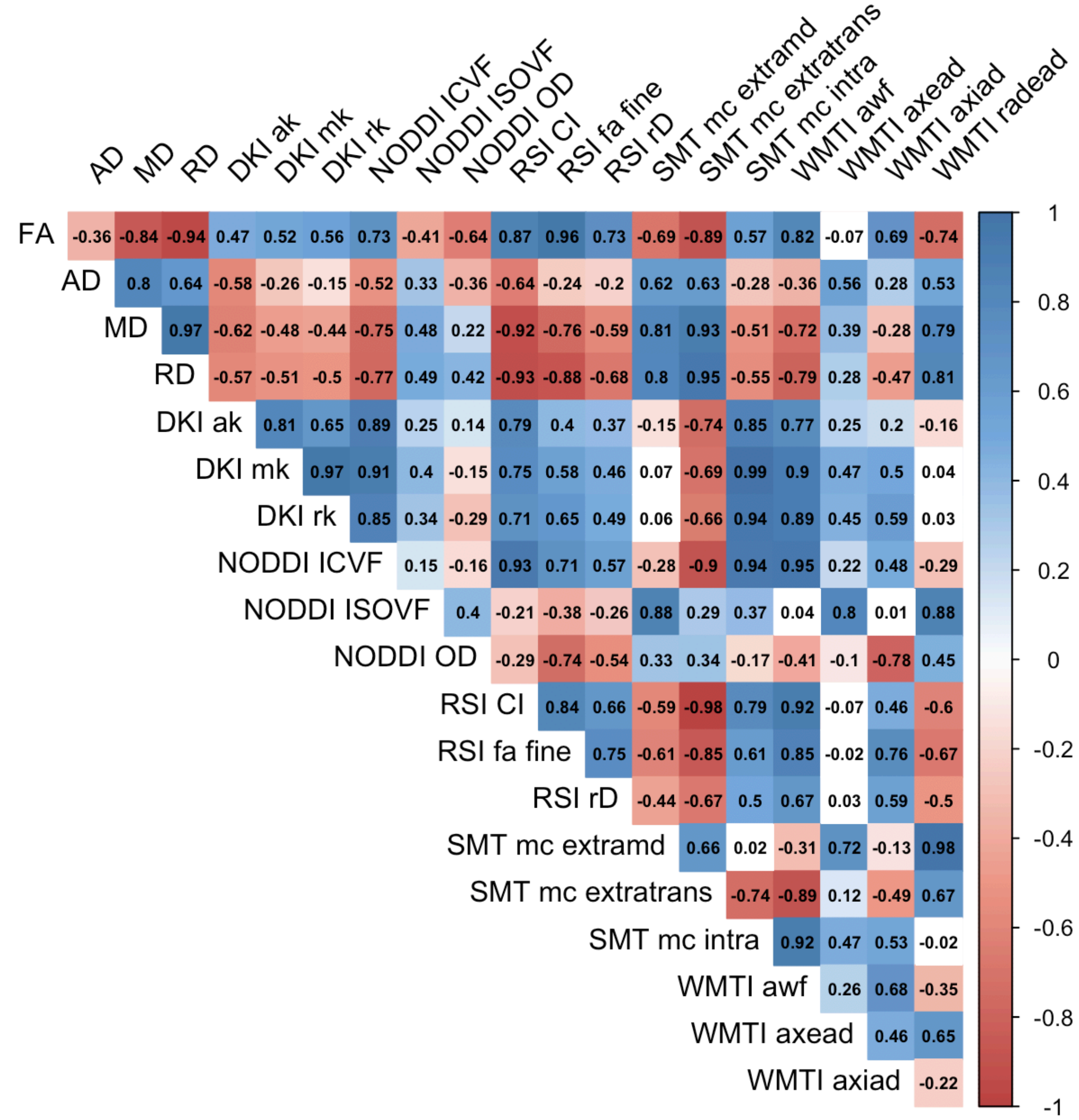


**SI Figure 2**. Correlation matrix showing the associations between all metrics (raw mean skeleton values).

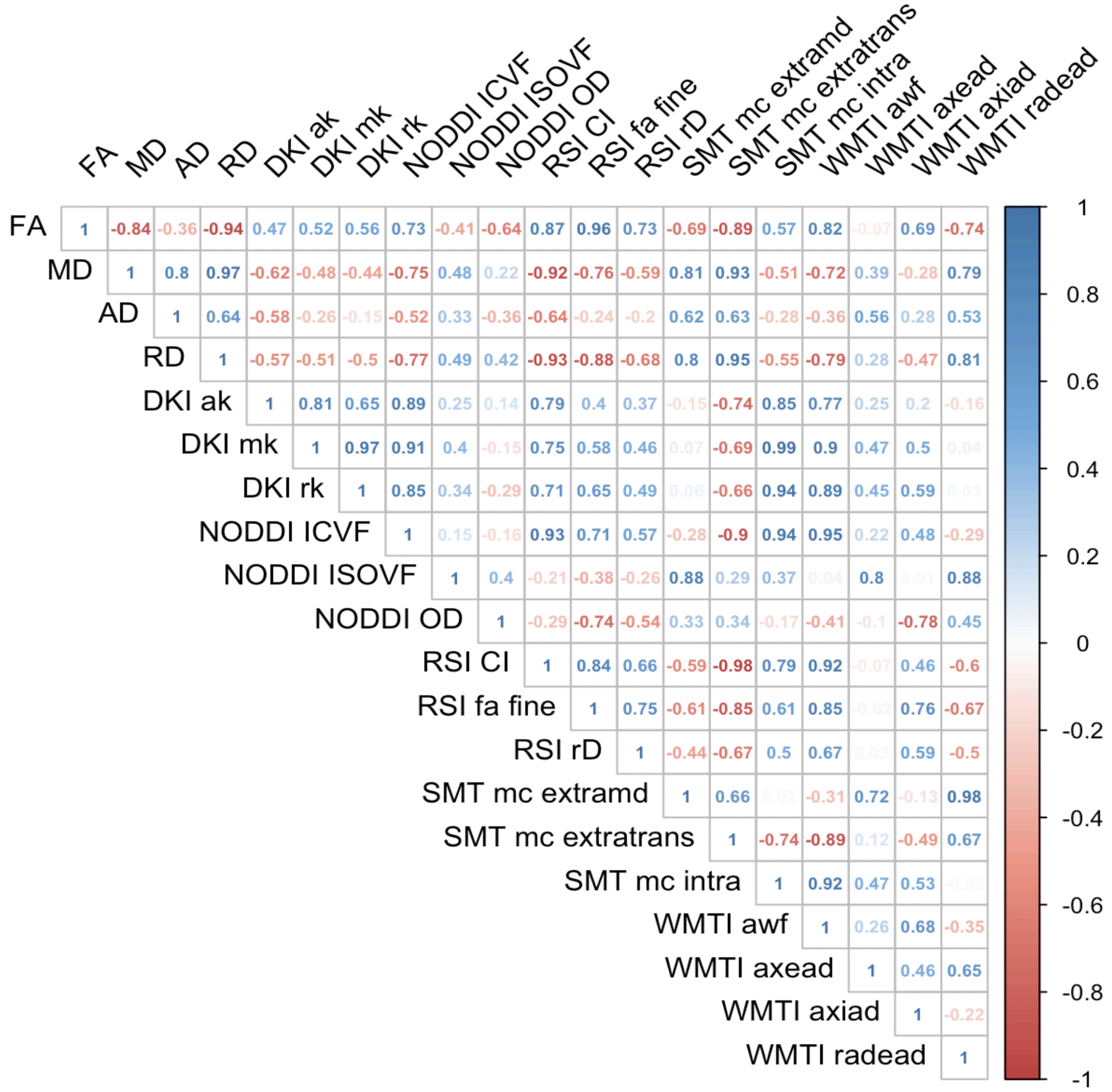


**SI Figure 3.** Correlation matrix showing the associations between all metrics using standardised values of the raw values.

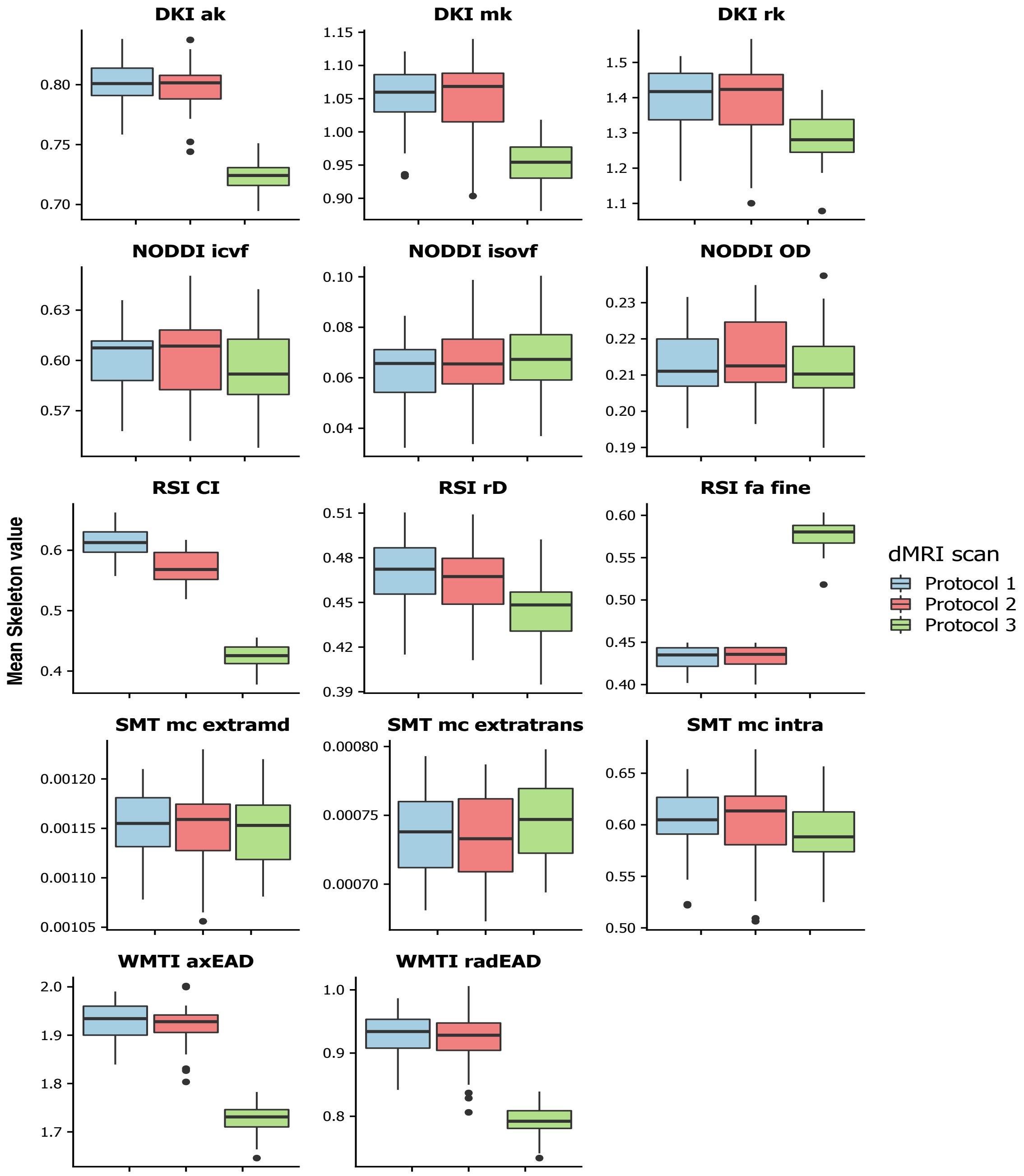


**SI Figure 4**: Showing Protocol 1: 60-30 directions for b1000/2000 s/mm$^2$. Protocol 2: 60-60 directions for b1000/2000 s/mm$^2$. Protocol 3: 60-60-60 directions with additional b3000 s/mm$^2$. Sample used in figure represents sub-sample of full study sample (n = 23 x 3 protocols).

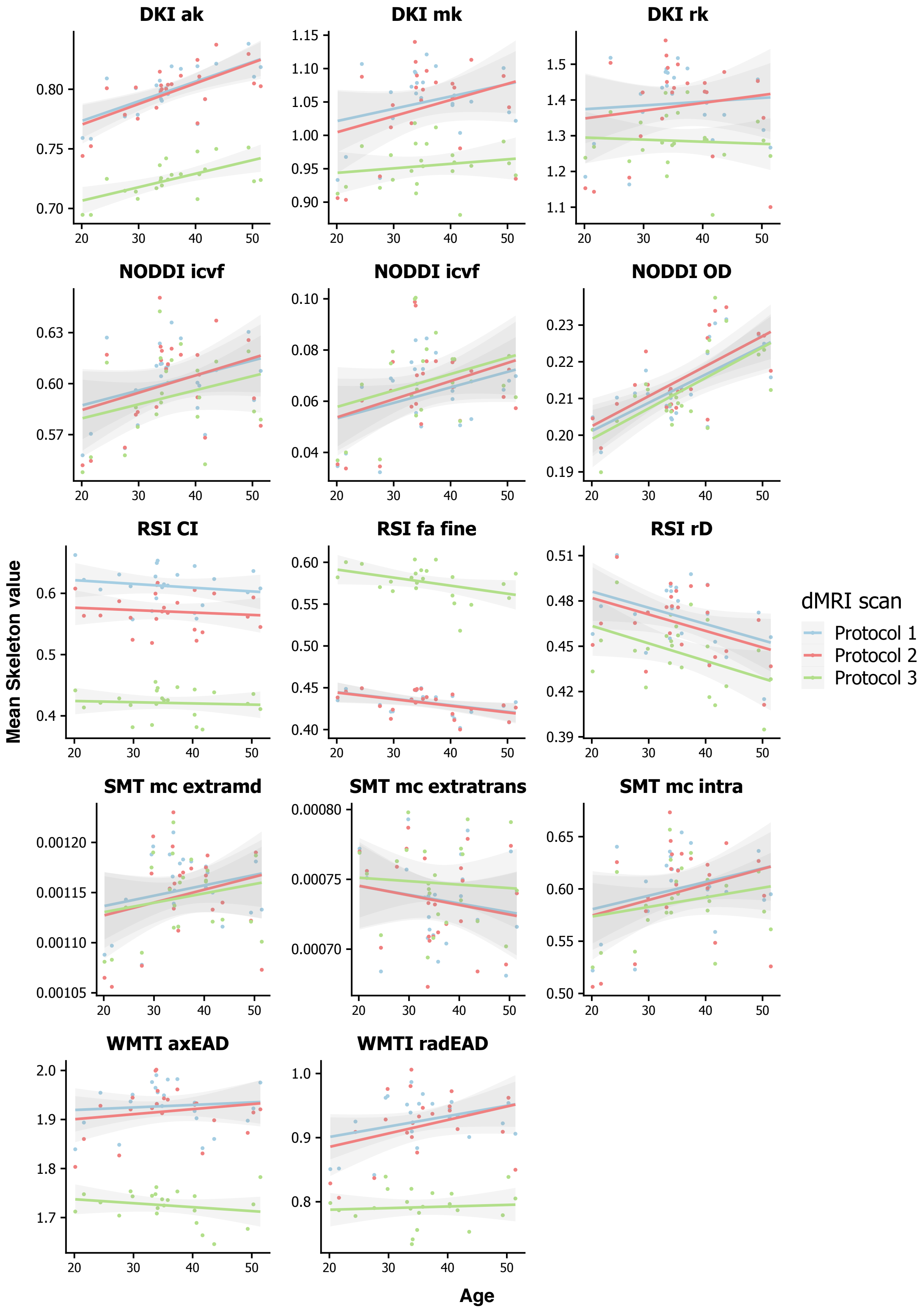


**SI Figure 5**. Showing Protocol 1: 60-30 directions for b1000/2000 s/mm$^2$. Protocol 2: 60-60 directions for b1000/2000 s/mm$^2$. Protocol 3: 60-60-60 directions with b3000 s/mm$^2$. Sample used in figure represents sub-sample of full study sample (n = 23 x 3 protocols).

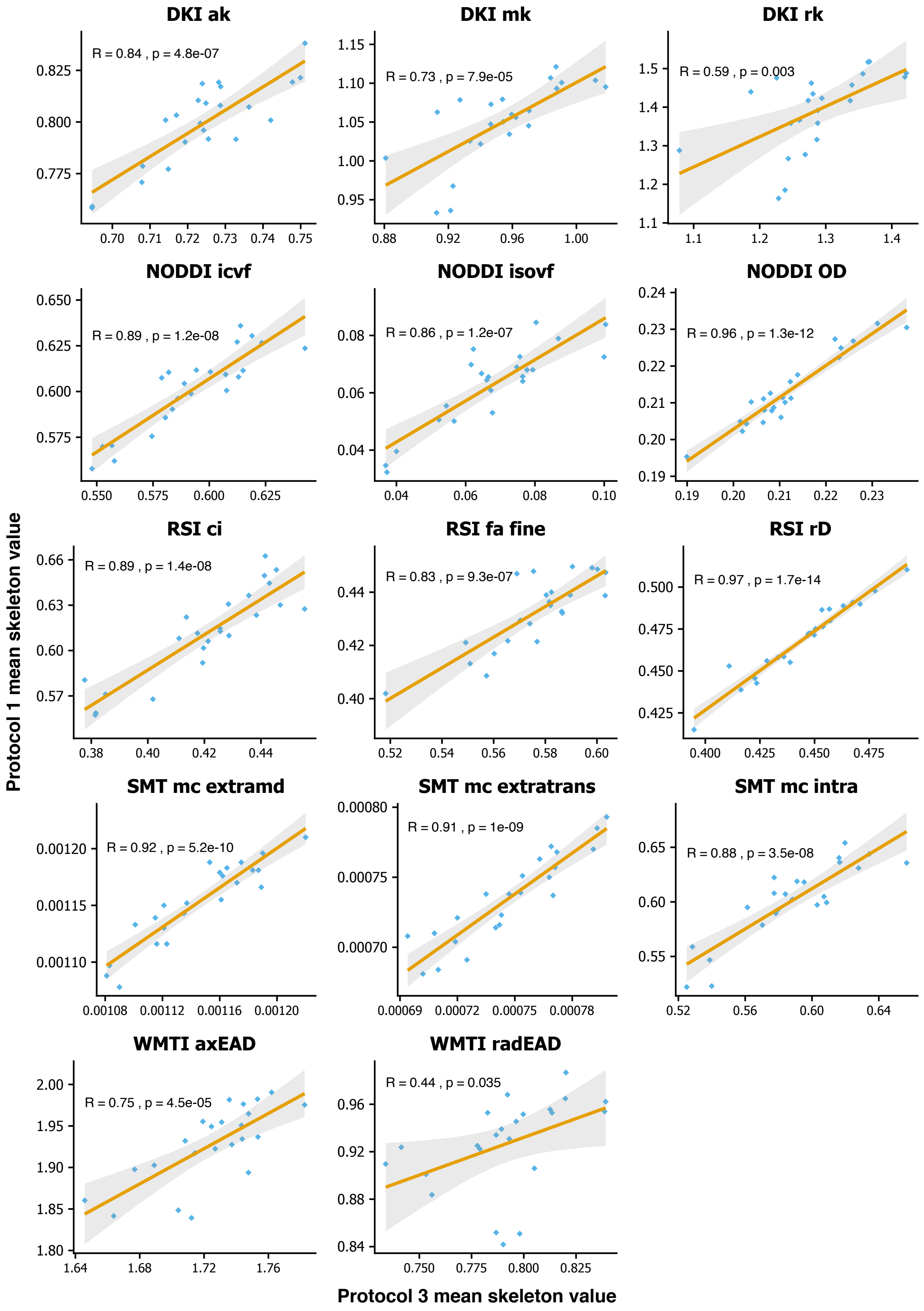


**SI Figure 6**. Showing Protocol 1: 60-30 directions with b1000/2000 s/mm$^2$. Protocol 2: 60-60 directions for b1000/2000 s/mm$^2$. Protocol 3: 60-60-60 directions with additional b3000 s/mm$^2$ values, where x axis is Protocol 1 factored by Protocol 3 (y axis).

**SI table 4.** Showing corrected *P* values ($p^{corr}$) corresponding to values displayed in SI Figure 6. Effect size estimated as R-squared ($R^2$).

| Diffusion metric | $p^{corr}$ | $R^2$ [95% CI] |
|---|---|---|
| DTI FA | 1.09 x $10^{-04}$ | 0.89 [0.57, 0.91] |
| DTI MD | 0.059 | 0.76 [0.23, 0.80] |
| DTI RD | 0.016 | 0.80 [0.31, 0.83] |
| DKI ak | 8.09 x $10^{-06}$ | 0.92 [0.66, 0.93] |
| DKI mk | 0.001 | 0.85 [0.45, 0.88] |
| DKI rk | 0.051 | 0.77 [0.24, 0.80] |
| NODDI ICVF | 2.01 x $10^{-07}$ | 0.94 [0.76, 0.95] |
| NODDI ISOVF | 1.99 x $10^{-06}$ | 0.93 [0.70, 0.94] |
| NODDI OD | 2.13 x $10^{-11}$ | 0.98 [0.90, 0.99] |
| RSI CI | 2.40 x $10^{-07}$ | 0.94 [0.75, 0.95] |
| RSI fa fine | 1.58 x $10^{-05}$ | 0.91 [0.64, 0.93] |
| RSI rD | 2.95 x $10^{-13}$ | 0.99 [0.93, 0.99] |
| SMT mc extramd | 8.81 x $10^{-09}$ | 0.96 [0.82, 0.97] |
| SMT mc extratrans | 1.73 x $10^{-08}$ | 0.96 [0.81, 0.96] |
| SMT mc intra | 6.03 x $10^{-07}$ | 0.94 [0.73, 0.95] |
| WMTI axEAD | 0.001 | 0.86 [0.48, 0.89] |
| WMTI radEAD | 0.597 | 0.66 [0.04, 0.72] |

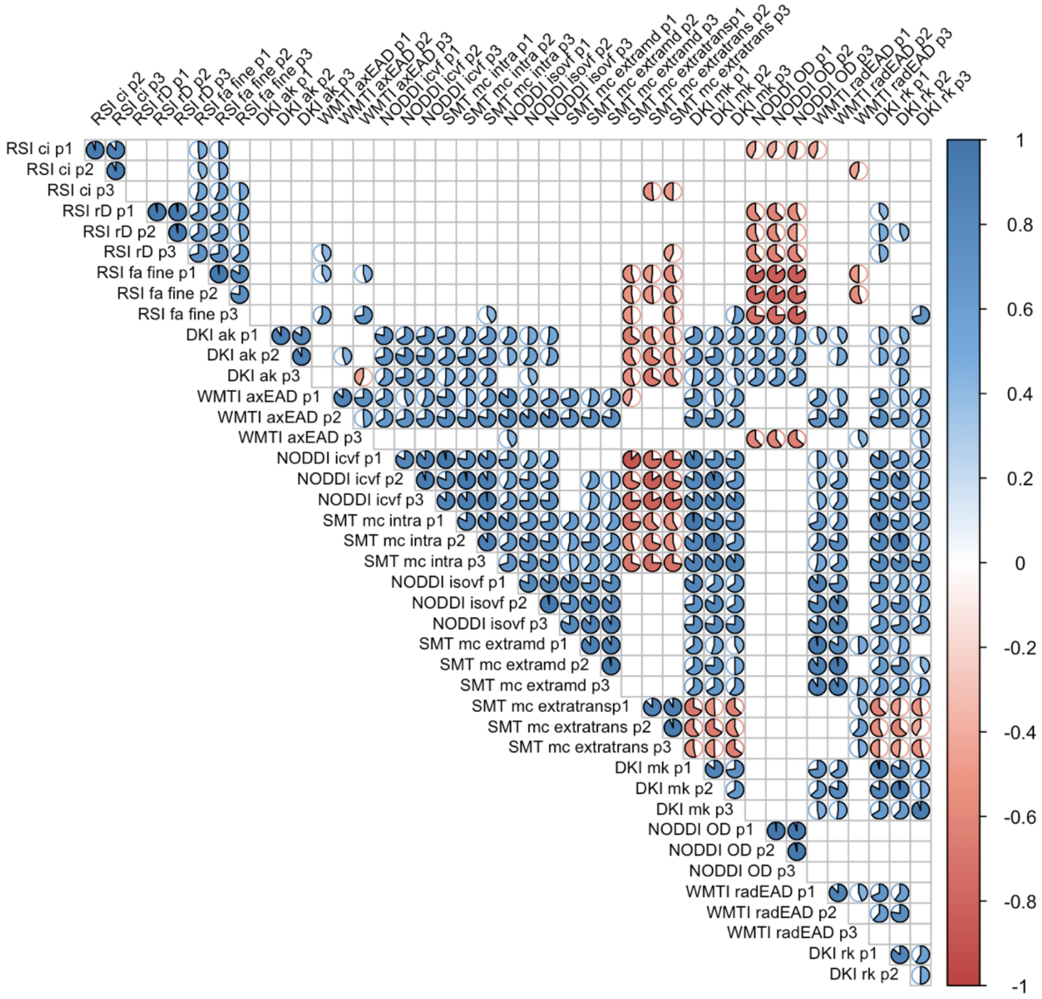


**SI Figure 7**. Correlation matrix showing associations between three dMRI protocols, where Protocol 1 = 60-30 directions (b1000/2000 s/mm$^2$), Protocol 2 = 60-60 directions (b1000/2000 s/mm$^2$), and Protocol 3 = 60-60-60 directions with additional b3000 s/mm$^2$. Non-significant boxes represented by blank boxes.

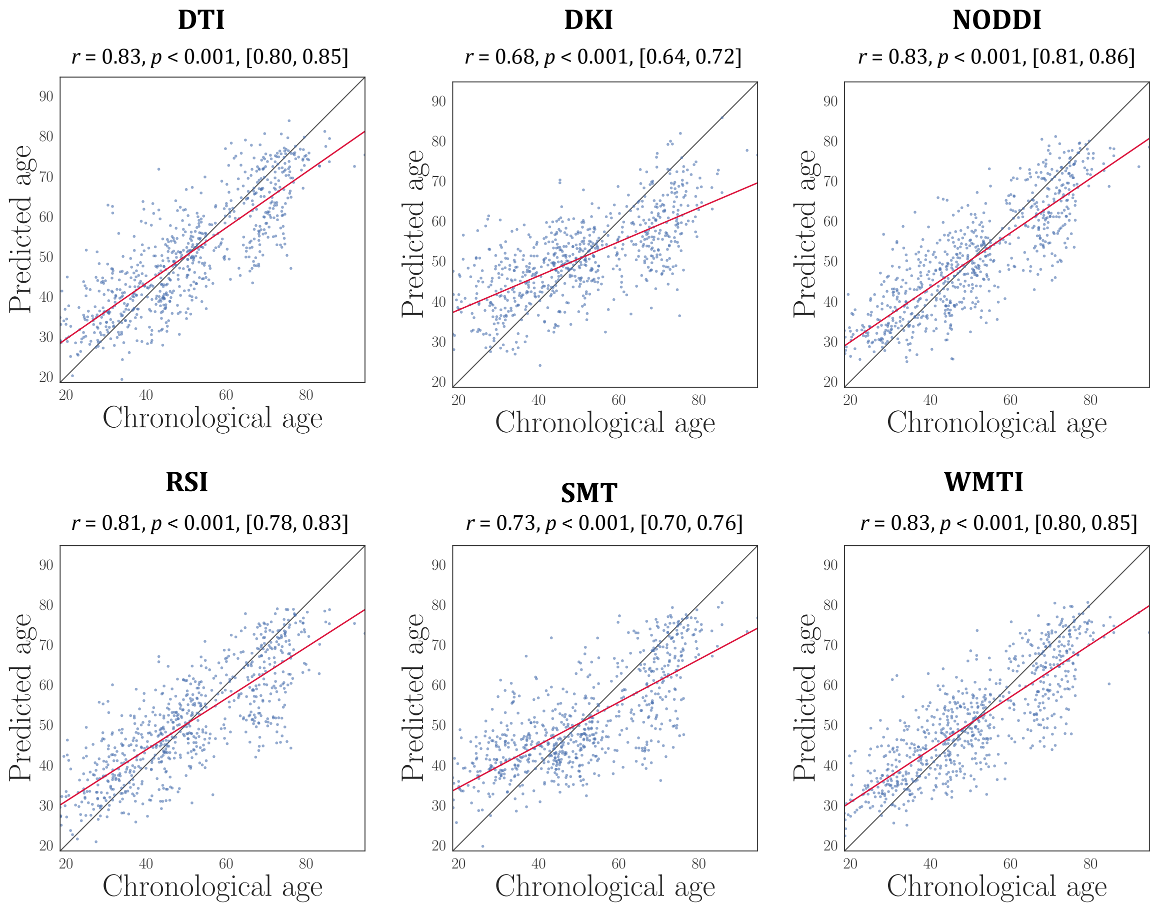


**SI Figure 8.** The association between predicted and chronological age shown for each of the modality-specific models. 95% confidence intervals are indicated in square brackets.

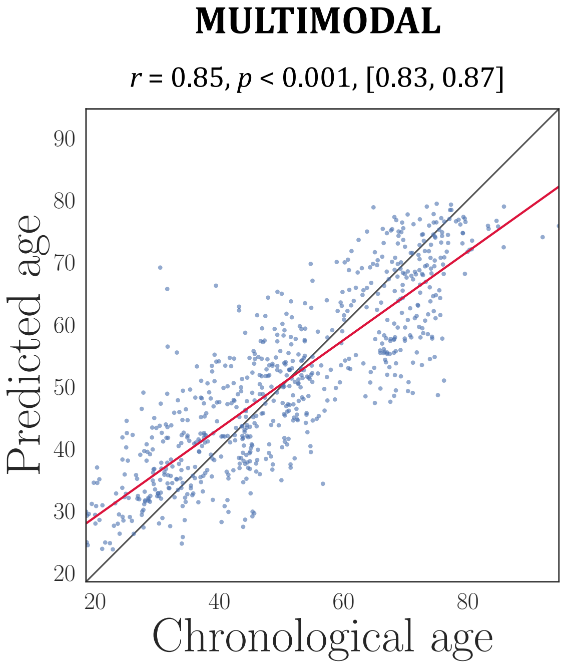


**SI Figure 9.** The association between predicted and chronological age for the multimodal model. 95% confidence intervals are indicated in square brackets.

**DTI FA**

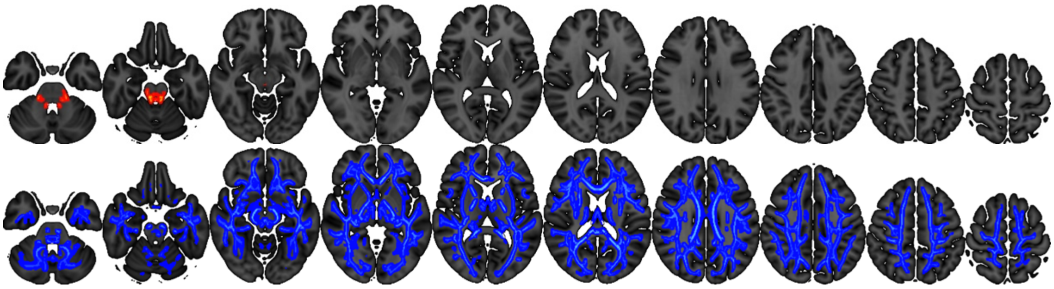

**DTI MD**

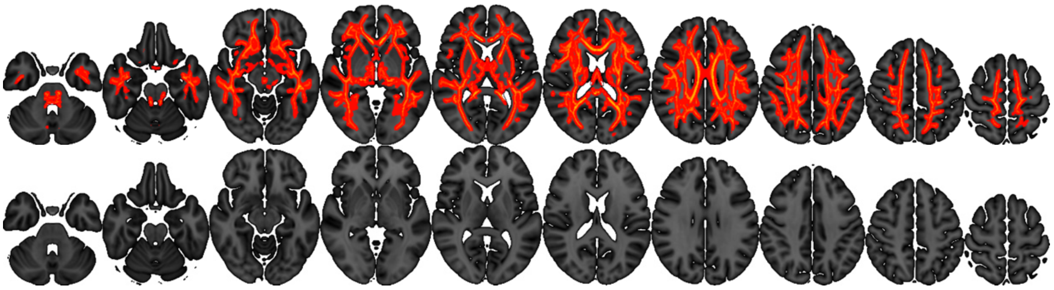

**DTI AD**

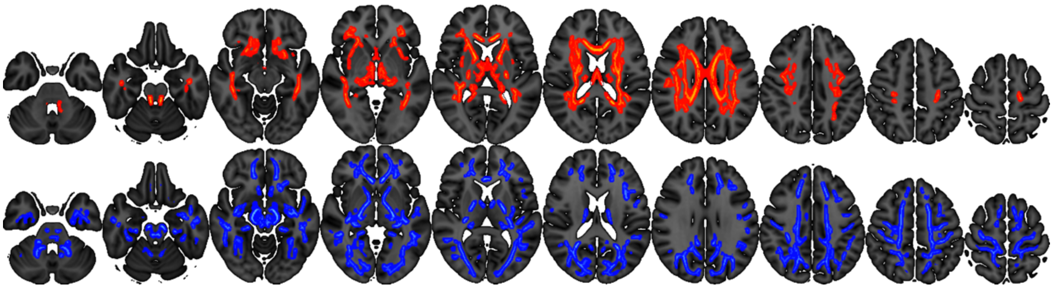

**DTI RD**

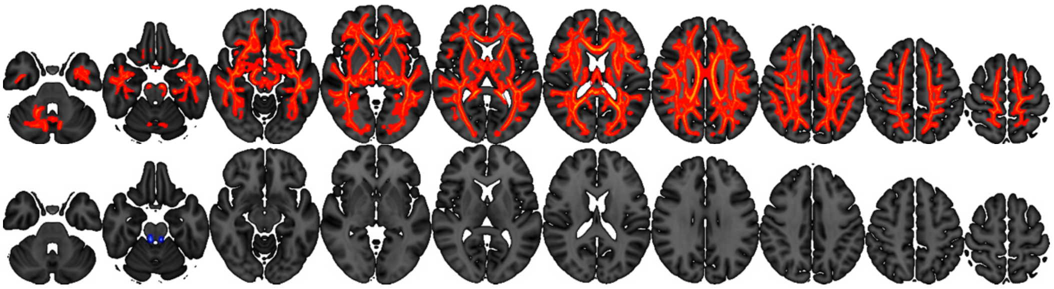

**DKI ak**

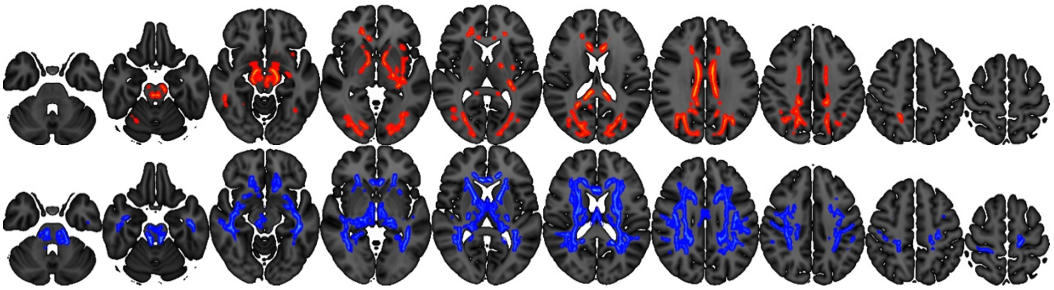

**DKI mk**

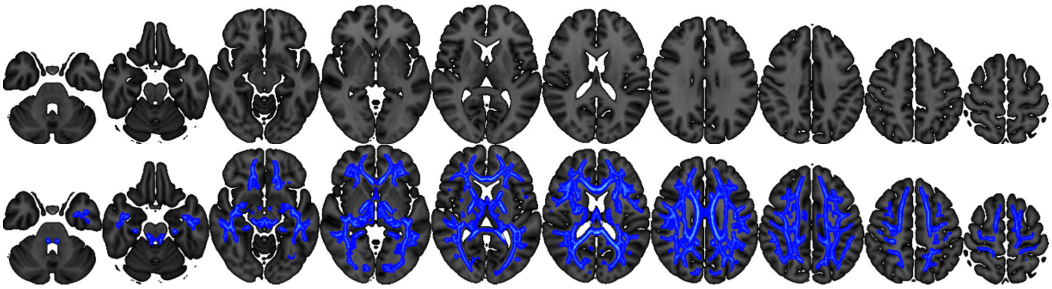

**DKI rk**

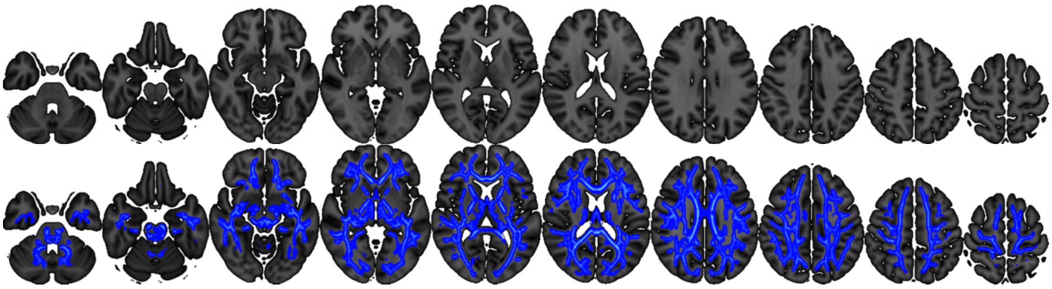

**NODDI icvf**

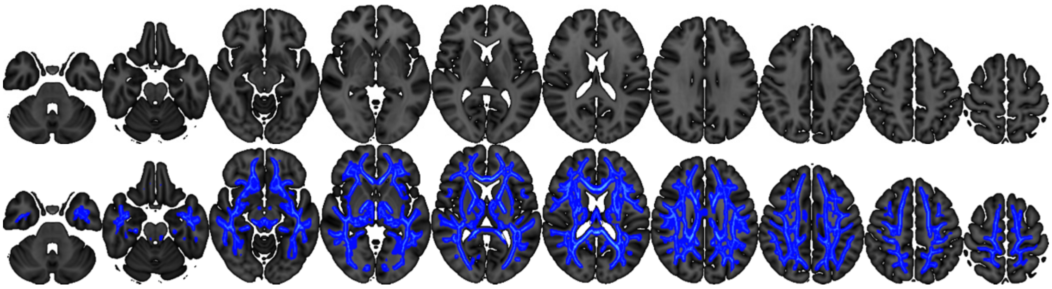

**NODDI isovf**

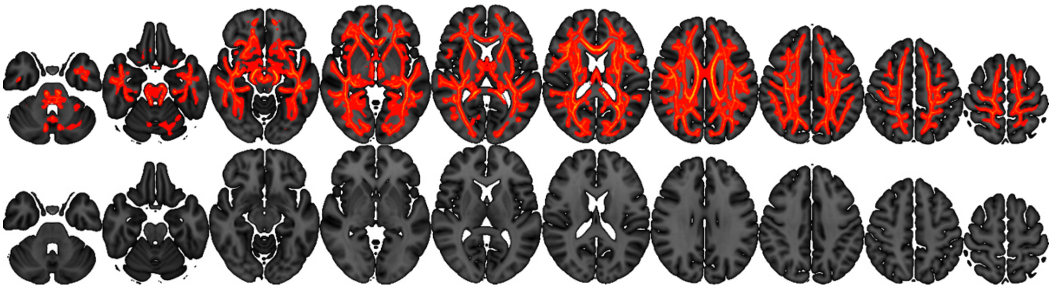

**NODDI od**

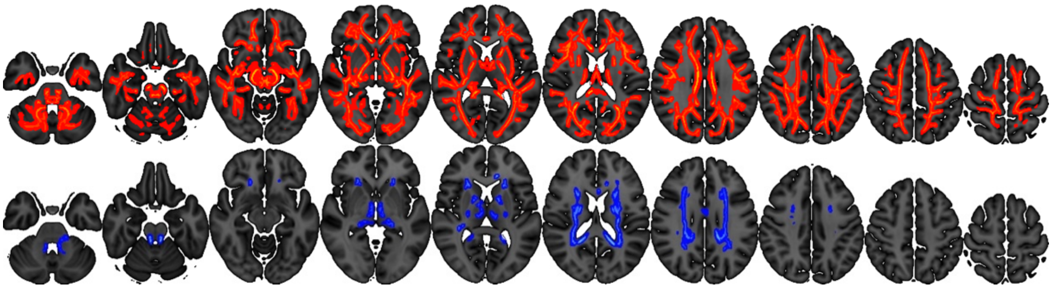

**RSI ci**

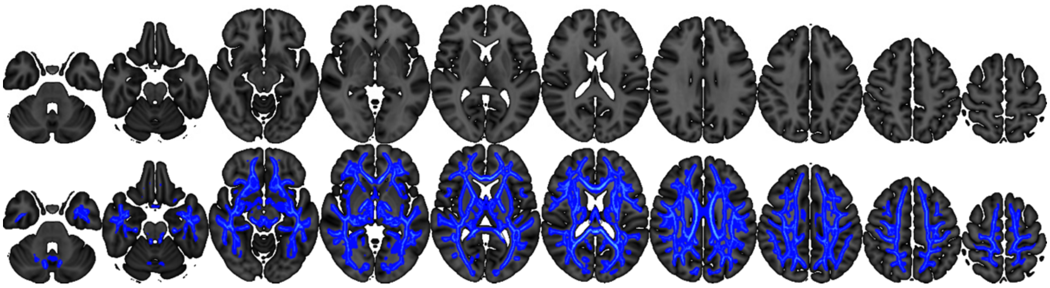

**RSI fa fine**

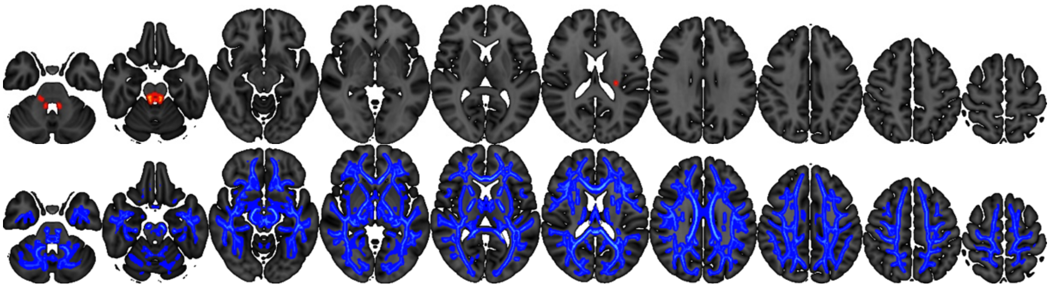

**RSI rD**

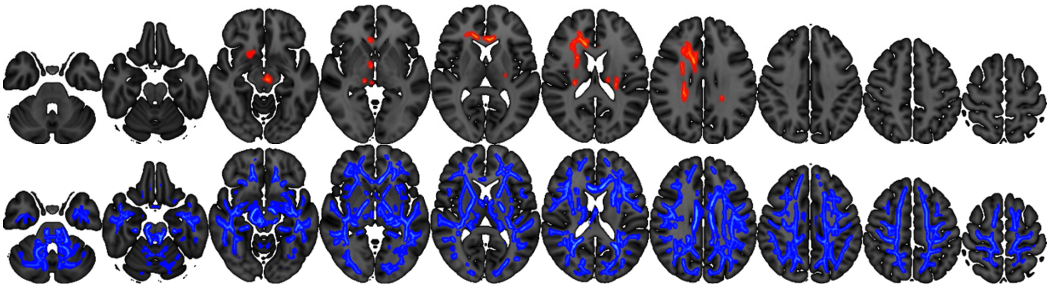

**SMT mc extramd**

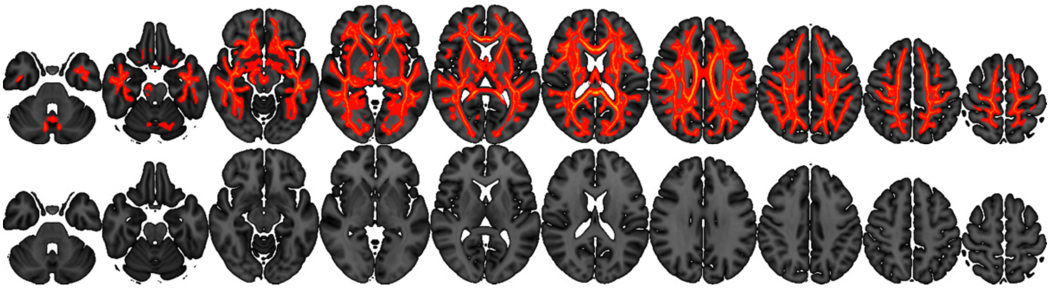

**SMT mc extratrans**

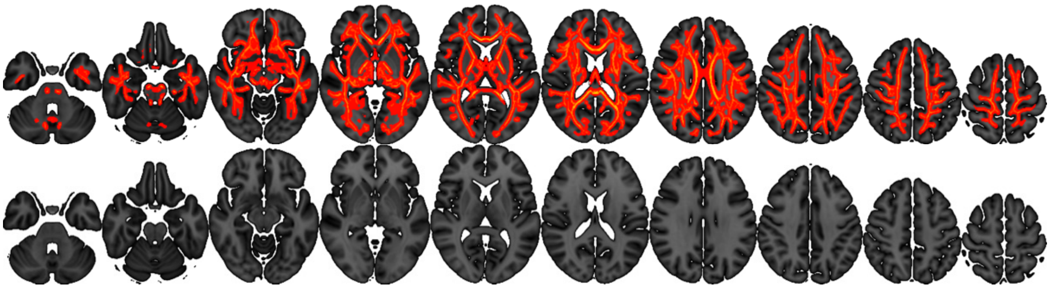

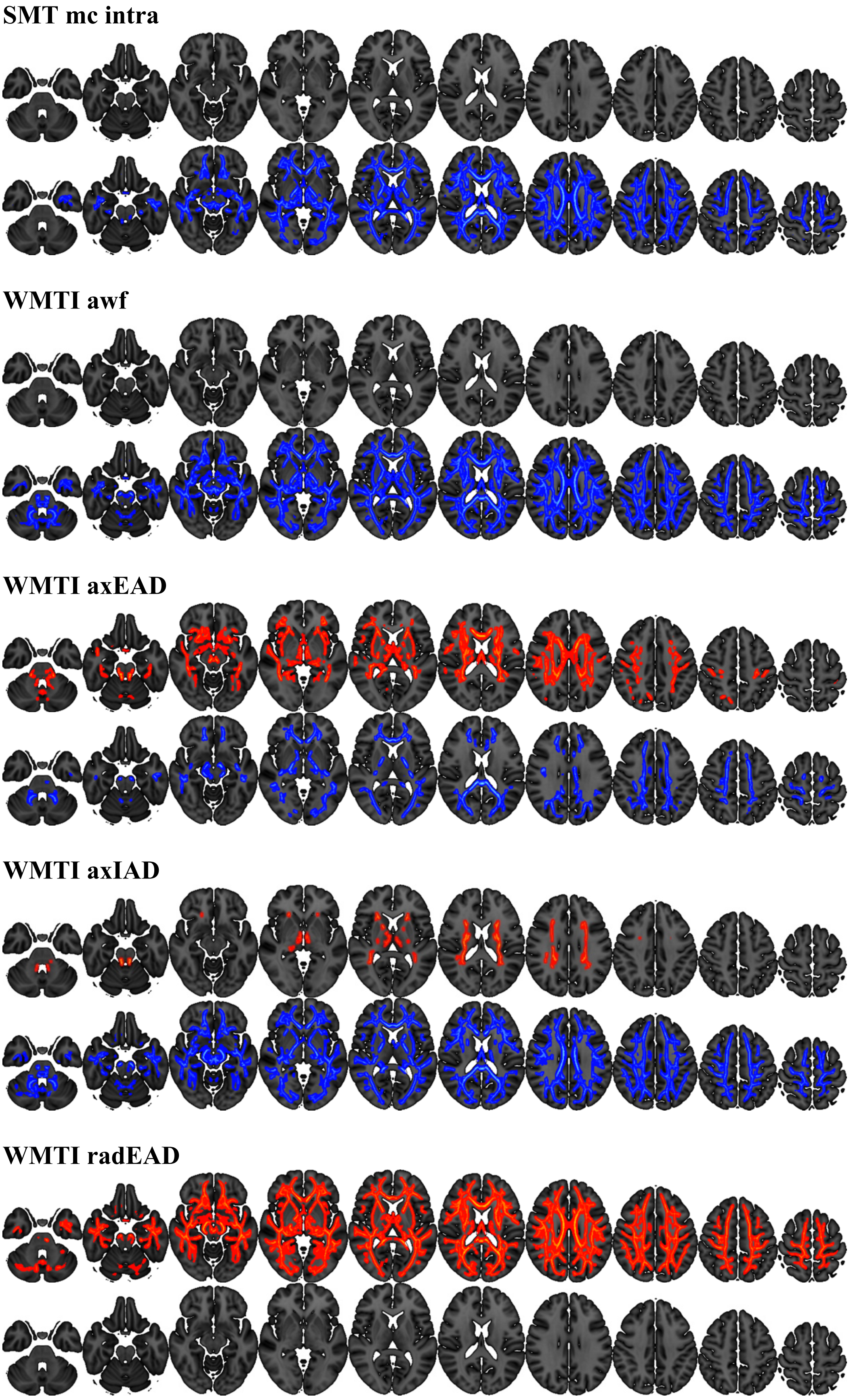


SI Figure 10: Voxel-wise analysis for each of the 21 metrics, corrected for multiple comparisons using permutation testing and TFCE. Voxels with one-tailed values of $p < 0.05$ are shown in red, displaying positive association with age, and in blue, displaying negative associations with age. From left to right, the following numbers represent the z-coordinate in MNI 1-mm space: 40, 50, 60, 70, 80, 90, 100, 110, 120, 130.
